# A fluorogenic chemically induced dimerization technology for controlling, imaging and sensing protein proximity

**DOI:** 10.1101/2023.01.04.522617

**Authors:** Sara Bottone, Zeyneb Vildan Cakil, Octave Joliot, Gaelle Boncompain, Franck Perez, Arnaud Gautier

## Abstract

Proximity between proteins plays an essential and ubiquitous role in many biological processes. Molecular tools enabling to control and observe the proximity of proteins are essential for studying the functional role of physical distance between two proteins. Here we present CATCHFIRE (Chemically Assisted Tethering of CHimera by Fluorogenic Induced REcognition), a chemically induced proximity technology with intrinsic fluorescence imaging and sensing capabilities. CATCHFIRE relies on genetic fusion to small dimerizing domains that interact upon addition of fluorogenic inducers of proximity that fluoresce upon formation of the ternary assembly, allowing real-time monitoring of the chemically induced proximity. CATCHFIRE is rapid and fully reversible, and allows the control and tracking of protein localization, protein trafficking, organelle transport and cellular processes, opening new avenues for studying or controlling biological processes with high spatiotemporal resolution. Its fluorogenic nature allowed furthermore the design of innovative biosensors for the study of various processes, such as signal transduction and apoptosis.

## INTRODUCTION

The many distinct activities of a cell are coordinated through the spatial and temporal organization of functionally interacting proteins via sequestration/translocation into specific compartments (e.g. organelles) or at the plasma membrane, or through the physical assembly of protein complexes via scaffold proteins. Technologies to chemically induce protein dimerization/proximity have been transformative for studying the role of spatiotemporal protein organization in various processes such as gene regulation, protein transport, signal transduction, metabolism, immune response, cell-cell communication^1^. Based on fusion to dimerization domains that interact in presence of chemical inducers of proximity, chemically induced proximity (CIP) enables to modulate protein interactions and thus cellular processes with high temporal control. Ideally, CIP should (i) rely on small dimerizing domains to avoid dysfunctional fusions, (ii) use non-toxic, easy to synthesize inducers of proximity, (iii) be rapid and reversible for dynamic studies, and (iv) have detection capabilities for real-time monitoring. Currently no technology meets all these criteria. CIP based on the rapamycin-inducible interaction of the FK506 binding protein (FKBP) and the FKBP-rapamycin binding domain (FRB) of the mammalian target of rapamycin (mTOR), is efficient and rapid, but is not reversible because of the high stability of the ternary assembly, and is limited by the biological activity of rapamycin^2,3^. These limits can be mitigated using non-toxic rapamycin analogs^4^ or alternative technologies based on dimerization domains involved in the sensing of plant hormones^5-8^, although these latter can suffer from rather bulky dimerizing domains.

In this study, we present a technology that addresses these limitations while incorporating unprecedented fluorescence imaging and sensing capabilities in its design. Currently, the monitoring of CIP requires fusion to fluorescent proteins (for colocalization studies or measurements of proximity by Förster resonance energy transfer^9^), or split reporters^10-12^. These additions inherently increase the risk of dysfunctional fusions and complicates the tagging of endogenous proteins by genome editing. In addition, colocalization or FRET studies require controlled stoichiometry. Split fluorescent reporters can circumvent this issue, however systems such as split fluorescent proteins^10^ suffer from slow and irreversible complementation, and, because of their irreversibility, can stabilize the assembly, preventing dynamic studies.

Our technology, dubbed CATCHFIRE (Chemically Assisted Tethering of CHimera by Fluorogenic Induced REcognition), relies on a small peptide of 11 amino acid residues dubbed ^FIRE^tag and a small protein domain of 114 amino acid residues dubbed ^FIRE^mate, whose interaction is reversibly mediated by fluorogenic inducers of proximity (called match) that fluoresce upon formation of the ternary assembly (**Fig. 1a,b**). The fluorogenic nature of CATCHFIRE allows real-time monitoring of proximity without the need for additional reporters. We showed that CATCHFIRE is rapid and reversible, and enables the control and tracking of cellular protein localization, intracellular protein trafficking, organelle positioning/transport and cellular processes, opening new avenues for studying and controlling biological processes with high spatiotemporal resolution. Moreover, we showed that the fluorogenic nature of CATCHFIRE allows the design of innovative biosensors for the detection of cellular activities.

**Figure 1.**
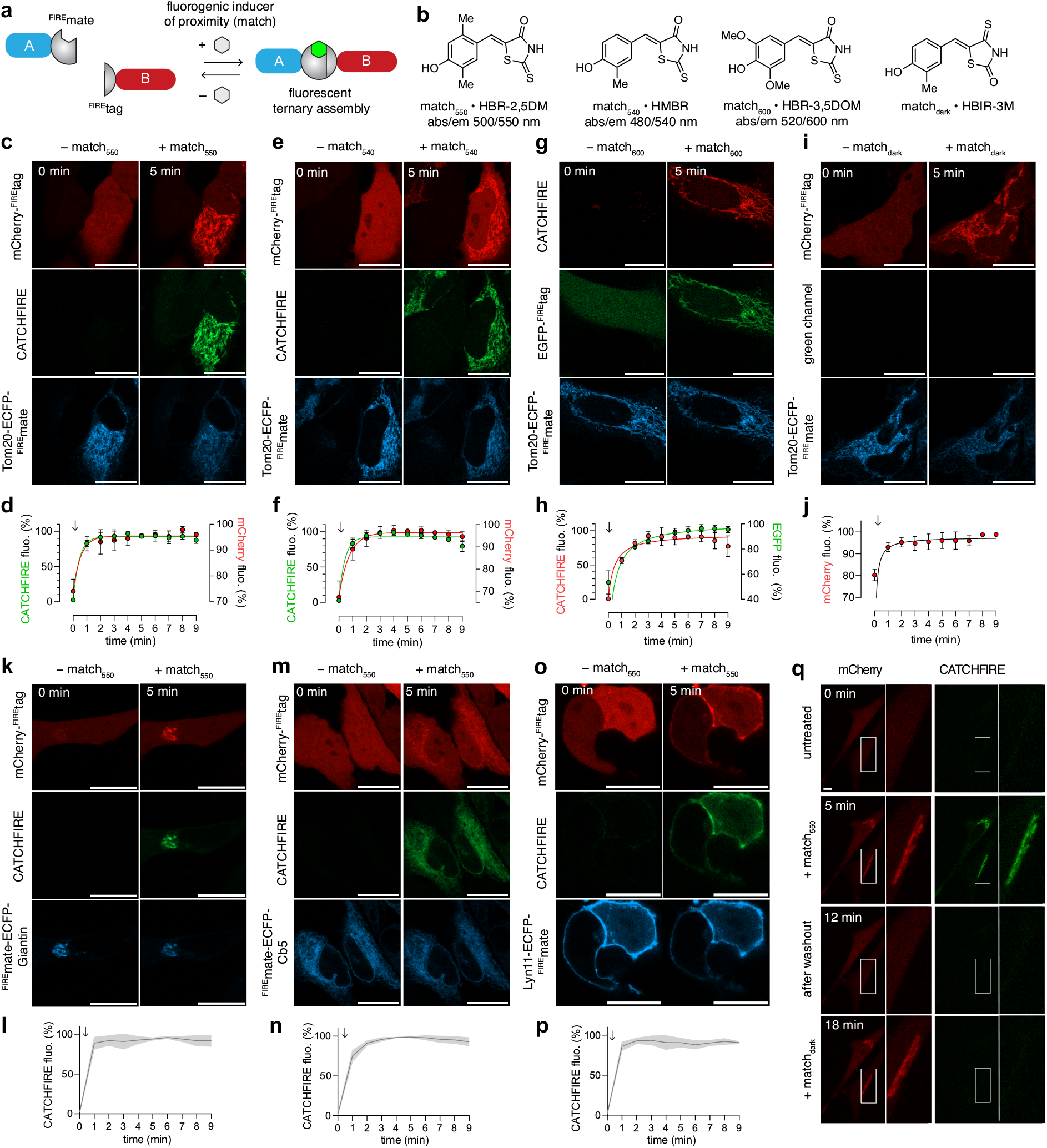
Chemically Activated Tethering of CHimera by Fluorogenic Induced REcognition (CATCHFIRE). **a** CATCHFIRE relies on the genetic fusion of two proteins A and B to ^FIRE^tag and ^FIRE^mate, two domains that can interact in a specific manner in presence of a fluorogenic inducer of proximity (called the *match*) that becomes fluorescent upon formation of the ternary assembly, enabling to induce the proximity of the proteins and visualize the recruitment process by fluorescence imaging. **b** Structures of the different inducers of proximity used in this study. **c-j** HeLa cells co-expressing mCherry-^FIRE^tag (**c-f**,**i**,**j**) or EGFP-^FIRE^tag (**g**,**h**) together with Tom20-ECFP-^FIRE^mate were treated with 10 µM of either match_550_ (**c**,**d**) or match_540_ (**e**,**f**) or match_600_ (**g**,**h**), or match_dark_ (**i**,**j**), and imaged by time-lapse confocal microscopy. **c**,**e**,**g**,**i** Representative confocal micrographs of cells before (0 min) and after (5 min) addition of the match (see **Movies S1, S6-S8**). Experiments were repeated three times with similar results. **d**,**f**,**h**,**j** Temporal evolution of the recruitment. Data represents the mean values ± SD of 20 cells (**d**), 10 cells (**f**), 10 cells (**h**), 10 cells (**j**) from three independent experiments. **k-p** HeLa cells co-expressing mCherry-^FIRE^tag and ^FIRE^mate-ECFP-Giantin (**k**,**l**), ^FIRE^mate-ECFP-Cb5 (**m**,**n**), Lyn11-^FIRE^mate-ECFP (**o**,**p**) were treated with 10 µM match_550_, and imaged by time-lapse confocal microscopy. **k**,**m**,**o** Representative confocal micrographs of cells before (0 min) and after (5 min) addition of match_550_ (see **Movies S9-S11**). Experiments were repeated three times with similar results. **l**,**n**,**p** Temporal evolution of the CATCHFIRE signal. Data represents the mean values ± SD of 10 cells (**l**), 17 cells (**n**), 22 cells (**p**) from three independent experiments. **q** HeLa cells co-expressing mCherry-^FIRE^tag and Giantin-ECFP-^FIRE^mate were treated with 10 µM match_550_, washed and then treated with 10 µM match_dark_. Cells were imaged by time-lapse spinning-disk microscopy. Representative micrographs of untreated cells, cells after addition of match_550_, cells after washout of match_550_, cells after addition of match_dark_. Experiments were repeated three times with similar results. Microscopy settings: **c-p** Red: ex/em 561/606-675 nm; Green: ex/em 488/508-570 nm; Cyan: ex/em 445/455-499 nm ; Scale bars are 20 µm; **q** Red: ex/em 561/604-664 nm; green: ex/em 488/525-545 nm. Scale bars are 10 µm.

## RESULTS

### CATCHFIRE design

^FIRE^tag and ^FIRE^mate were generated by bisection of the fluorescent chemogenetic reporter pFAST^13^. pFAST is a promiscuous and improved version of FAST (Fluorescence-Activating and absorption-Shifiting Tag), a 14 kDa protein that binds and stabilizes the fluorescent state of fluorogenic chromophores. Non-fluorescent when free, these so-called fluorogens enable to label fusion proteins with high contrast without the need for fluorogen washout^14^. FAST was previously used to design a split fluorescent reporter for detecting protein-protein interactions through bisection at position 114-115. Because the two complementary fragments of FAST exhibit low complementation in presence of fluorogen but can fully complement when in close proximity, they allow the detection of interacting proteins^11^. When applying the same splitting strategy on pFAST, we discovered that pFAST_1-114_ (dubbed ^FIRE^mate) and pFAST_115-125_ (dubbed ^FIRE^tag) interact in presence of fluorogens, allowing the development of a fluorogenic recruitment system that we called CATCHFIRE. We showed that fluorogens can act as molecular glue and force the assembly of ^FIRE^mate and ^FIRE^tag by fusing the two fragments to the C-termini of FKBP and FRB. Evaluation by flow cytometry of the complementation efficacy in presence of fluorogens in HEK293T cells treated or not with rapamycin (to induce or not FKBP and FRB interaction), revealed that the ternary fluorescent assembly forms efficiently regardless of the presence of rapamycin (**Fig. S1**), demonstrating that fluorogens (hereafter called ‘matches’) can induce the interaction between ^FIRE^tag and ^FIRE^mate.

To evaluate whether ^FIRE^mate and ^FIRE^tag self-complement in absence of match, we spatially separated the two fragments. We fused ^FIRE^tag to the C-terminus of the red fluorescent protein mCherry to produce a diffuse cytosolic expression, and ^FIRE^mate to the C-terminus of the N-terminal domain of the mitochondrial outer membrane protein TOM20 (TOM20_1-34_), so that ^FIRE^mate is immobilized, facing the cytosol. An enhanced cyan fluorescent protein (ECFP) was additionally inserted to monitor the localization of the ^FIRE^mate fusion. In untreated cells, mCherry-^FIRE^tag showed diffused cytosolic localization while co-expressed TOM20-ECFP-^FIRE^mate localized at mitochondria, suggesting that the two fragments do not spontaneously complement at the expression levels explored (**Figure 1c,d**). Treatment with match_550_ (a.k.a. HBR-2,5DM) triggered rapid translocation of the red fluorescent mCherry fusion from the cytosol to the mitochondria surface and led to the simultaneous appearance of a strong green-yellow fluorescent signal at the mitochondria (**Figure 1c,d** and **Movie S1**). Match_550_-induced proximity was further evidenced by a decrease of the mitochondrial ECFP signal, in agreement with Förster resonance energy transfer (FRET) between ECFP and the ternary fluorescent assembly (**Fig. S2**). These experiments showed that match_550_ could induce the proximity of proteins fused to ^FIRE^mate and ^FIRE^tag respectively, while lighting up the recruitment process. Fast time-lapse confocal microscopy showed that the half-time of recruitment was 20 seconds, suggesting that CATCHFIRE is rapid and mainly limited by cellular uptake and diffusion of the match (**Fig. S3a**,**b** and **Movie S2**). Long-term time-lapse microscopy showed that the fluorogenic recruitment was stable over long period of time (**Figure S3c**,**d** and **Movie S3**). Fusion of ^FIRE^tag at the N-terminus of mCherry, or insertion of ^FIRE^tag in beween two mCherry further demonstrated the efficiency of the approach regardless of the position of ^FIRE^tag (**Figure S4** and **Movies S4-S5**).

### CATCHFIRE enables imaging with various colors

Match_550_ induces the formation of a green-yellow fluorescent ternary assembly. As chromophores with various spectral properties can bind pFAST^13^, we next explored their use as (fluorogenic) chemical inducer of proximity to extend the ‘colors’ available for detecting CATCHFIRE. We showed that the green fluorescent match_540_ (a.k.a. HMBR) and the red fluorescent match_600_ (a.k.a. HBR-3,5DOM) enabled efficient fluorogenic induction of proximity (**Figure 1e-h** and **Movies S6-S7**). The non-fluorogenic match_dark_ (a.k.a. HBIR-3M) was also shown to induce efficient recruitment (**Figure 1i,j** and **Movie S8**), which can be interesting when rapid recruitment, but no fluorescence induction, is needed. The ability to induce proximity without generating fluorescence could find various applications in biological and medical research as demonstrated by conventional CIP tools.

### Control and tracking of protein localization

To demonstrate the versatility of CATCHFIRE, we next anchored ^FIRE^mate at the membrane of various organelles or cellular structures so that it faced the cytosol. We fused the transmembrane domain (3131-3259 amino acid residues) of Giantin^15^, or the transmembrane domain (100-134 amino acid residues) of the cytochrome b5 (Cb5) to the C-terminus of ^FIRE^mate to target it to the Golgi apparatus and the endoplasmic reticulum respectively. We also fused ^FIRE^mate at the C-terminus of the Lyn11 inner membrane targeting sequence. For each fusion, an ECFP was inserted to monitor the localization of the fusion. Each fusion was co-expressed together with cytosolic mCherry-^FIRE^tag. Addition of match_550_ or match_540_ resulted in rapid and specific rerouting of the red fluorescent mCherry-^FIRE^tag to the organelle expressing the ^FIRE^mate and the simultaneous appearance of green-yellow fluorescence on the respective organelle (**Fig. 1k-p, Fig. S5, Movies S9-S14**), demonstrating efficient fluorogenic induced recruitment. The rerouting of mCherry-^FIRE^tag occurs with comparable kinetics regardless of the organelle. CATCHFIRE can thus be used to induce fast fluorogenic recruitment of proteins to sub-cellular compartments.

### CATCHFIRE allows reversible interaction

Next, we showed that CATCHFIRE is reversible and that a simple washout of the fluorogenic inducer of proximity is sufficient to abolish the interaction. We expressed mCherry-^FIRE^tag and ^FIRE^mate-ECFP-Giantin, and induced fluorogenic recruitment by adding match_550_ to the cells. Washout of match_550_ led to simultaneous release of mCherry fusion into the cytosol and disappearance of the green fluorescence, in agreement with the rapid dissociation of the fluorescent ternary assembly, demonstrating the full reversibility of the recruitment (**Figure 1q, Movie S15**). Recruitment could be induced again by addition of the non-fluorescent match_dark_, showing the possibility to induce repeated catch and release cycles (**Figure 1q, Movie S15**). The ability to reversibly control protein proximity on-demand opens great prospect to study cellular processes with high temporal resolution.

### CATCHFIRE combination with other CIP technologies

We next asked whether CATCHFIRE could be combined with the FRB-FKBP-rapamycin CIP system in order to control two interactions. We expressed the chimeric fusion mCherry-FKBP-^FIRE^tag together with both FRB-ECFP-Giantin and TOM20- ^FIRE^mate, so that mCherry-FKBP-^FIRE^tag was in excess. Sequential addition of, first, match_550_ and, then rapamycin, allowed the rerouting of mCherry-FKBP-^FIRE^tag first to mitochondria (through ^FIRE^tag-^FIRE^mate interaction) and then to the Golgi apparatus (through FKBP-FRB interaction) (**Fig. S6**), demonstrating that the two systems are orthogonal and can be used for controlling two interactions in the same experiment.

### CATCHFIRE set-up without basal activity

Concerned that high expression level could lead to some degree of self-association of ^FIRE^tag and ^FIRE^mate in absence of match, we developed a caged version of ^FIRE^tag to prevent any interaction with ^FIRE^mate in case one needs to ensure very low match-independent interaction. To do so, we took advantage of the photoactive yellow protein (PYP), pFAST’s ancestral protein^13,14^, which cannot bind any pFAST chromophores.

Fusion of ^FIRE^tag to the N-terminal domain of the apo PYP (PYP_2-114_) resulted in a protein (caged^FIRE^tag) that folds like a circular permutation of PYP because of the intramolecular interaction between ^FIRE^tag and PYP_1-114_ (**Fig. S7** and **S8a**), masking thus ^FIRE^mate. Analysis of the complementation efficacy of FKBP-caged^FIRE^tag and FRB-^FIRE^mate in presence and absence of rapamycin at various match_550_ concentrations by flow cytometry allowed us to show that caged^FIRE^tag could still complement with ^FIRE^mate in presence of match_550_, regardless of the proximity of the two domains (**Fig. S8b**). We observed that, in agreement with the caging of ^FIRE^tag, higher concentrations of match_550_ were however needed to reach full complementation in absence of rapamycin. We propose that, in presence of match_550_, the formation of a more stable ternary assembly provides the driving force to uncage ^FIRE^tag for efficient association with ^FIRE^mate via a mutually exclusive folding mechanism. Next, we expressed mCherry-caged^FIRE^tag together with ^FIRE^mate-ECFP-Giantin in HeLa cells. In absence of match_550_, the two proteins did not colocalized, confirming that caged^FIRE^tag and ^FIRE^mate exhibit no significant binding affinity (**Fig. S8c**,**d**). Treatment with match_550_ resulted in efficient, albeit slower, recruitment of mCherry-caged^FIRE^tag to the Golgi, demonstrating efficient complementation of the two domains (**Fig. S8c**,**d, Movie S16)**.

### Control and tracking of nucleocytoplasmic protein trafficking

Next, we showed that CATCHFIRE could allow the development of systems to induce protein motility such as nucleocytoplasmic protein trafficking. Nucleocytoplasmic trafficking is tightly regulated through mechanisms typically involving nuclear localization signals (NLS) and nuclear export signal (NES). CATCHFIRE-mediated nuclear protein export was demonstrated using NLS-mCherry-^FIRE^mate as a NLS-containing cargo and NES-ECFP- ^FIRE^mate as an export NES-containing partner. NES-ECFP- ^FIRE^mate is mainly localized in the cytosol, but can transiently traffic in the nucleus by diffusion (**Fig. 2a**). Treatment with match_550_ resulted in rapid nuclear export of NLS-mCherry-^FIRE^tag through interaction with NES-ECFP- ^FIRE^mate, as judged by a nuclear-to-cytoplasmic shift of mCherry fluorescence and the appearance of cytoplasmic green fluorescence (**Fig. 2a-c** and **Movie S17**). This result confirmed that in protein assemblies containing both NLS and NES, the export activity prevails over the import activity^16^.

**Figure 2.**
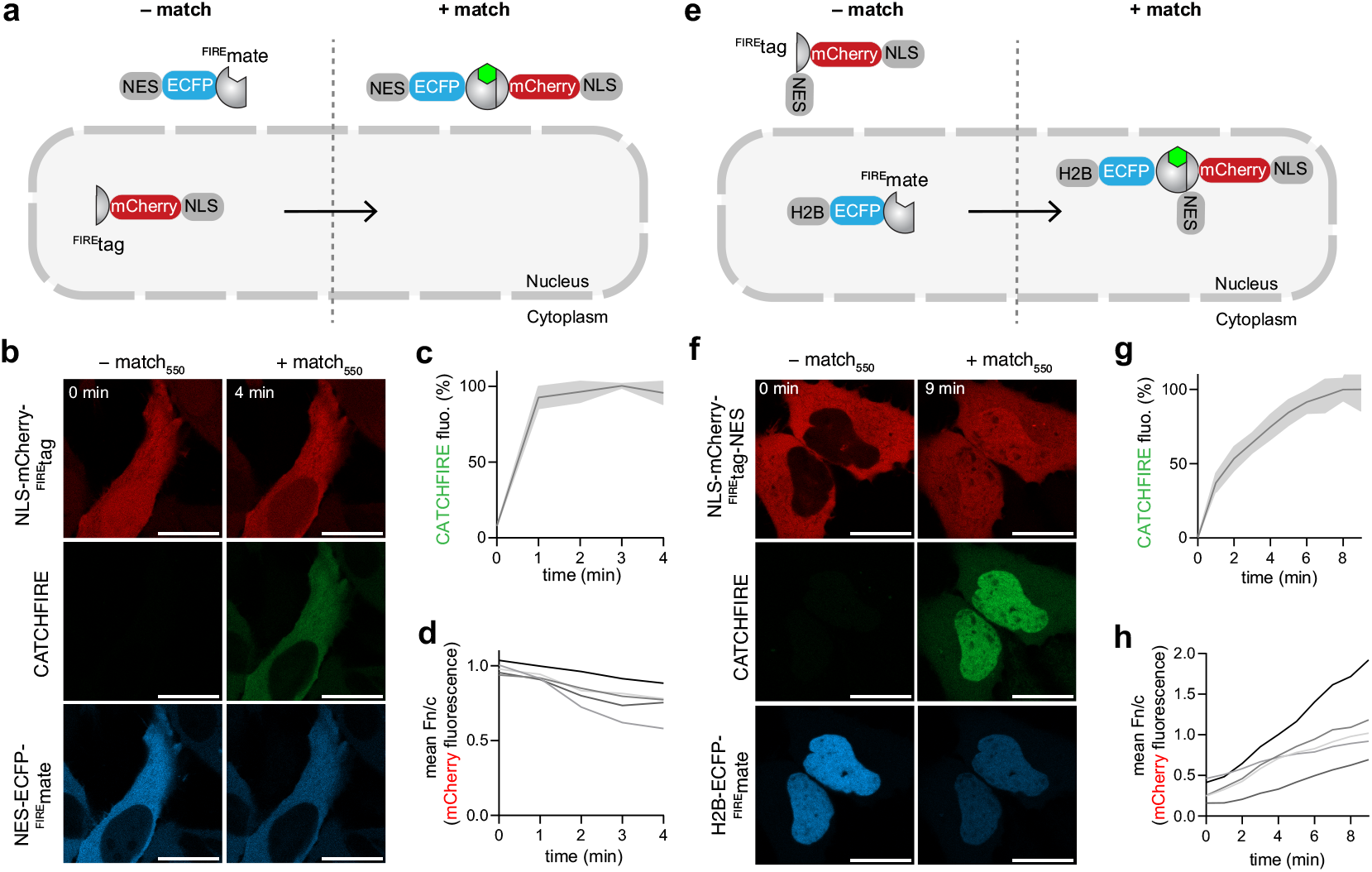
Control and tracking of nucleocytoplasmic shuttling. **a-d** HeLa cells co-expressing NLS-mCherry-^FIRE^tag and NES-ECFP- ^FIRE^mate were treated with 10 µM match_550_ and imaged by time-lapse confocal microscopy. **b** Representative confocal micrographs before and after treatment with match_550_ (see also **Movie S17**). Experiments were repeated three times with similar results. **c** Temporal evolution of the CATCHFIRE signal. Data represent the mean ± SD of n = 5 cells from two independent experiments. **d** Temporal evolution of the ratio nucleus-to-cytoplasm mCherry fluorescence of n = 5 cells from 2 independent experiments. **e-h** HeLa cells co-expressing NLS-mCherry-^FIRE^tag-NES and H2B-ECFP- ^FIRE^mate were treated with 10 µM match_550_ and imaged by time-lapse confocal microscopy. **f** Representative confocal micrographs before and after treatment with match_550_ (see also **Movie S18**). Experiments were repeated three times with similar results. **g** Temporal evolution of the CATCHFIRE signal. Data represent the mean ± SD of n = 5 cells from two independent experiments. **f** Temporal evolution of the ratio nucleus-to-cytoplasm mCherry fluorescence of n = 5 cells from 2 independent experiments. **b**,**f** Red: ex/em 561/606-675 nm; green: ex/em 488/508-570 nm; cyan: ex/em 445/455-499 nm. Scale bars are 20 µm.

Conversely, CATCHFIRE enabled to mediate nuclear accumulation of a cytosolic cargo containing a NES and NLS (NLS-mCherry-^FIRE^tag-NES). H2B-ECFP- ^FIRE^mate was used here to trap NES-mCherry-^FIRE^tag-NLS transiently trafficking to the nucleus. Treatment with match_550_ allowed to sustainably trap NES-mCherry-^FIRE^tag-NLS transiently trafficking in the nucleus, inducing overall a cytosol-to-nucleus localization shift through a sink effect (**Fig. 2d-f** and **Movie S18**).

Overall this set of experiments suggests that the CATCHFIRE assay could be suitable to control and track protein trafficking between different cell compartments.

### Control and tracking of secretory protein traffic

As CATCHFIRE allows reversible interaction, we tested its use for the Retention Using Selective Hooks (RUSH) assay^17^ to control and track the traffic of secretory proteins. The RUSH system allows the synchronization of secretory protein traffic through the expression of two proteins: the hook, which is stably expressed in a donor compartment, and the reporter, which can reversibly interact with the hook. Upon reversion of the interaction, the reporter is released and traffics to its final compartment. We thus created an endoplasmic reticulum (ER) hook by fusing ^FIRE^mate with a C-terminal ER retention signal (Lys-Asp-Glu-Leu; KDEL), and three reporter proteins by fusing either the tumor necrosis factor-α (TNF) or the α-mannosidase II (ManII) to the N-terminus of a red fluorescent protein (mCherry or mApple)-^FIRE^tag, or a glycosylphosphatidylinositol (GPI) anchor to the C-terminus of mApple-^FIRE^tag (**Fig. 3**). To ensure efficient retention of the reporter by the hook, cells were treated after transfection with match_550_ for 24 h. At steady-state, TNF-mCherry-^FIRE^tag, mApple- ^FIRE^tag-GPI and ManII-mApple-^FIRE^tag were localized in the ER (**Fig. 3b-d**). To release the reporters, match_550_ was washed out, leading to a rapid loss of the green fluorescence and fast ER export. After 10 min, the three reporters were detected in the Golgi (**Fig. 3b-d** and **Movies S19-S21**). For TNF-mCherry-^FIRE^tag, granule-like post-Golgi carriers appeared after 15 min (**Fig. 3b, Movie S19**), while plasma membrane expression of mApple-^FIRE^tag-GPI was visible after 60 min (**Fig. 3c, Movie S20**). Interestingly, in the case of ManII-mApple-^FIRE^tag, KDEL-mediated retrograde traffic from the Golgi apparatus to the ER could be induced by re-addition of match_550_ (**Fig. 3d, Movie S21**). This set of experiments demonstrated the ability of CATCHFIRE to be used to synchronize and control the trafficking of secretory proteins. Compared to the conventional RUSH system that uses tetrameric streptavidin as a hook for the retention of reporters fused to streptavidin-binding peptides, and addition of biotin for the release, our CATCHFIRE-based RUSH system, in addition to bringing intrinsic fluorescence to monitor cargo release, benefits from the use of a monomeric hook, ^FIRE^mate, and from being reversible, opening new prospects for studying the mechanisms of trafficking.

**Figure 3.**
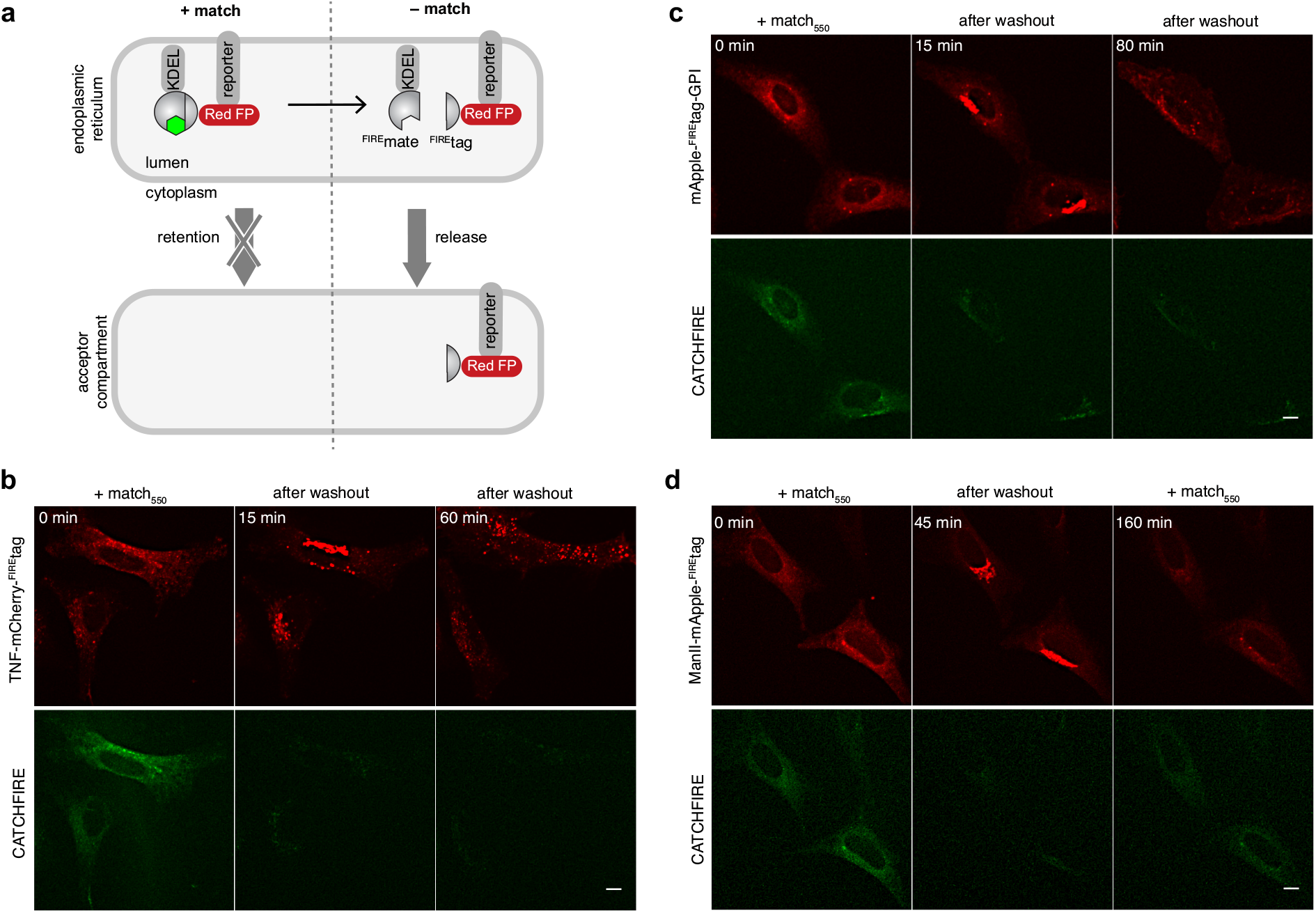
Control and tracking of secretory protein trafficking. **a** Schematic illustrating how ^FIRE^mate-KDEL acts as a hook to retain ^FIRE^tag-ged reporter in the endoplasmic reticulum in presence of the match. Release is induced by washing of the match, allowing trafficking of the reporter to its acceptor compartment. **b** HeLa cells co-expressing ^FIRE^mate-KDEL and TNF-mCherry-^FIRE^tag were treated with match_550_ for 24 h and imaged by spinning-disk microscopy after washout of match_550_. Representative micrographs before washout, 15 min and 60 min after washout (see also **Movie S19**). Experiments were repeated three times with similar results. **c** HeLa cells co-expressing ^FIRE^mate-KDEL and mApple- ^FIRE^tag-GPI were treated with match_550_ for 24 h and image by spinning-disk microscopy after washout of match_550_. Representative micrographs before washout, 15 min and 80 min after washout (see also **Movie S20**). Experiments were repeated three times with similar results. **c** HeLa cells co-expressing ^FIRE^mate-KDEL and ManII-mApple-^FIRE^tag were treated with match_550_ for 24 h and image by spinning-disk microscopy after washout of match_550_. Match_550_ was re-added after 50 min. Experiments were done in 25 µg/mL cycloheximide. Representative micrographs before washout, 45 min after washout and at 160 min (110 min after re-addition of match_550_) (see also **Movie S21**). Experiments were repeated four times with similar results. Red: ex/em 561/604-664 nm; green: ex/em 488/525-545 nm. Scale bars are 10 µm. All images were enhanced by Denoise.ai on NIS-Elements Ver.5.42 and by 2D-deconvolution on NIS-Elements Ver.5.42.

### Control and tracking of organelle positioning

CATCHFIRE can efficiently control the movement of proteins between compartments, and we wondered whether it could be used to alter and track the positioning of organelles through recruitment of specific molecular motors. We focused on lysosomes, whose motility and positioning are essential in various cellular processes, such as destruction of pathogens, plasma membrane repair, cell adhesion and migration, tumor invasion and metastasis, apoptosis, metabolism and gene regulation^18,19^. ^FIRE^tag was targeted to lysosomes using a lysosomal-associated membrane protein (LAMP)1-mCherry-^FIRE^tag fusion, and ^FIRE^mate was fused to the N-terminus of the motor domain of kinesin-like KIF17^20^, a member of the kinesin-2 family of plus-end directed microtubule-based motor proteins. Addition of match_550_ induced movement of the lysosomes from the perinuclear region to the cell periphery, in agreement with an induced anterograde transport. Lysosomes were repositioned within 30-45 minutes (**Fig. 4, Movie S22**). Washout of match_550_ led to repositioning of the lysosome toward the perinuclear region. Directionality of the movement could be reversed a second time by re-addition of match_550_. This experiment demonstrated that CATCHFIRE could efficiently control organelle positioning opening exciting prospects for getting insights into the functional effects of organelle displacement or to alter intracellular organization.

**Figure 4.**
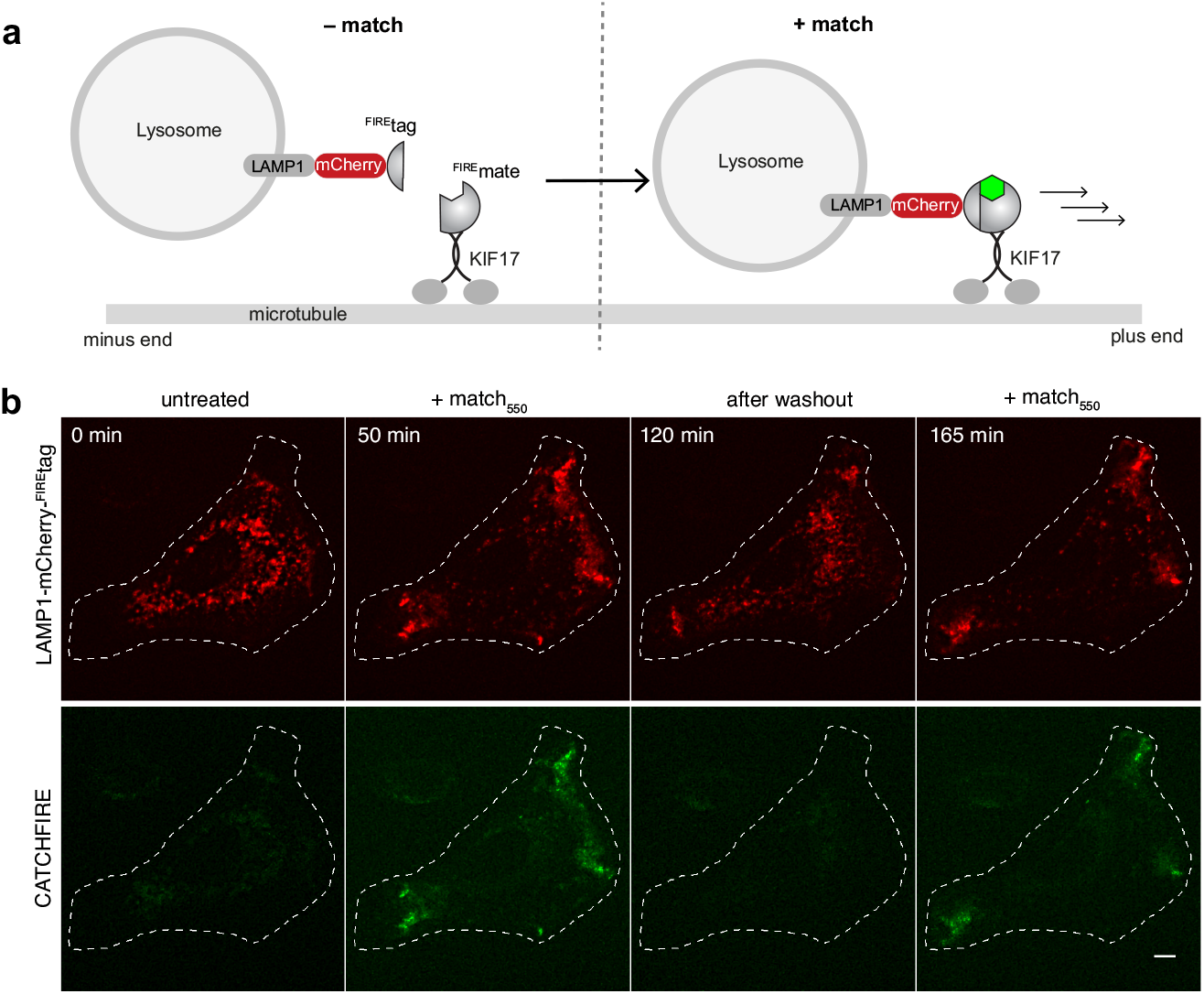
Control and tracking of organelle positioning. **a** Schematic illustrating how match-induced interaction between LAMP1-mCherry-^FIRE^tag and ^FIRE^mate-KIF17 allows the chemically induced anterograde transport of lysosomes. **b** HeLa cells co-expressing LAMP1-mCherry-^FIRE^tag and ^FIRE^mate-KIF17 were treated with match_550_ and imaged by spinning-disk microscopy. After 60 min, match_550_ was washed out and cells were imaged for 60 min without match_550_, before re-addition of match_550_ for 60 min. Representative micrographs of each step (see also **Movie S22**). Experiments were repeated four times with similar results. Red: ex/em 561/604-664 nm; green: ex/em 488/525-545 nm. Scale bars are 10 µm. Images were enhanced by Denoise.ai on NIS-Elements Ver.5.42 and by 2D-deconvolution on NIS-Elements Ver.5.42.

### Control of mitophagy

As spatiotemporal control of protein localization can allow to tune cell behavior, we explore the use of CATCHFIRE to control key cellular mechanisms. We focused on mitophagy, the selective degradation of mitochondria by autophagy. One of the main pathways involves the protein kinase PINK1^21^. In unhealthy cells, depolarization of the mitochondrial membrane impairs PINK1 internalization and leads to its accumulation at the mitochondrial outer membrane, initiating mitophagy. We investigated CATCHFIRE-mediated mitophagy through translocation of PINK1 at the mitochondrial outer membrane using PINK1-mCherry-^FIRE^tag. A fusion ^FIRE^mate-ECFP-(mitochondrial fission 1 protein)Fis1 was generated as mitochondrial outer membrane anchor. Addition of match_550_ resulted in rapid translocation of PINK1-mCherry-^FIRE^tag to the mitochondrial outer membrane (**Fig. S9** and **Movie S23**). Observations after 2h of treatment revealed strong morphology change of mitochondria in agreement with the triggering of mitophagy (**Fig. 5** and **Fig.S10**), showing that recruitment of PINK1 at the mitochondrial outer membrane was sufficient to initiate mitophagy. This experiment showed how CATCHFIRE could be used to control cellular processes through recruitment of specific effectors.

**Figure 5.**
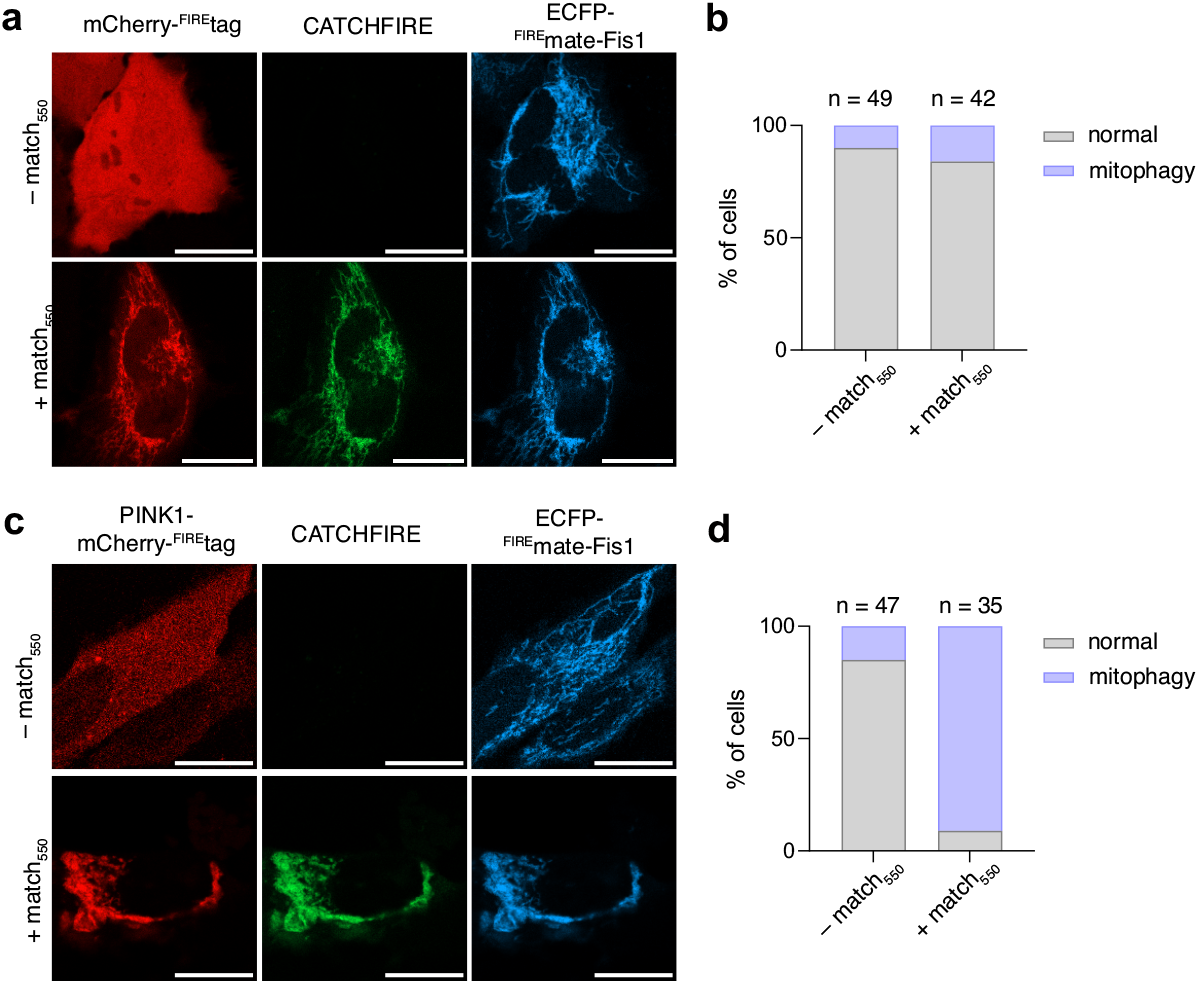
Fluorogenic induced recruitment of PINK1 to mitochondria induces mitophagy. **a-d** HeLa cells co-expressing ECFP-^FIRE^mate-Fis1 and either **a**,**b** mCherry-^FIRE^tag (negative control) or **c**,**d** PINK1-mCherry-^FIRE^tag were treated without or with 10 µM match_550_ for 2 h. **a**,**c** Representative micrographs (see also **Fig. S10**). Experiments were repeated three times with similar results. Red: ex/em 561/606-675 nm; green: ex/em 488/508-570 nm; cyan: ex/em 445/455-499 nm. Scale bars are 20 µm. **b**,**d** Quantification of the percentage of cells with normal morphology (grey) or mitophagy-induced morphology (purple). Number (*n*) of analyzed cells are reported.

### Design of fluorescent sensors for monitoring cellular activity

The fluorogenic nature of CATCHFIRE provides unprecedented opportunities for the design of sensors that switch on upon changes in cellular activity. Activation of signaling pathways or cellular processes often results in change of localization of one or several proteins. As CATCHFIRE requires ^FIRE^tag and ^FIRE^mate to be in the same cellular compartment to occur, it is possible to design sensors that switch on upon activation of a specific pathways by stably anchoring ^FIRE^mate within a specific organelle and tracking a reporter fused to ^FIRE^tag. To demonstrate the ability to design such fluorescent sensors, we first designed a sensor of signaling cascade activation. We used p65, a transcription factor of the NF-κB family involved in cell proliferation, apoptosis, cytokine production, and oncogenesis^22^. In most unstimulated cells, p65 is sequestered in the cytoplasm by IκB inhibitory proteins. Stimulation of cells leads to degradation of IκB, and allows the translocation of p65 to the nucleus, resulting in expression of target genes^22^. A sensor of p65 nuclear import was designed using p65-mCherry-^FIRE^tag as reporter and H2B-ECFP-^FIRE^mate as a nuclear anchor. In unstimulated cells, no nuclear green fluorescence was observed upon treatment with match_550_ in agreement with p65-mCherry-^FIRE^tag being sequestered in the cytoplasm, preventing thus efficient fluorogenic induced interaction with nuclear H2B-ECFP-^FIRE^mate (**Fig. 6a-c**). Cell stimulation led to nuclear CATCHFIRE, in agreement with nuclear translocation of p65-mCherry-^FIRE^tag (**Fig. 6d-f**), demonstrating that CATCHFIRE can be used for sensing signal transduction.

**Figure 6.**
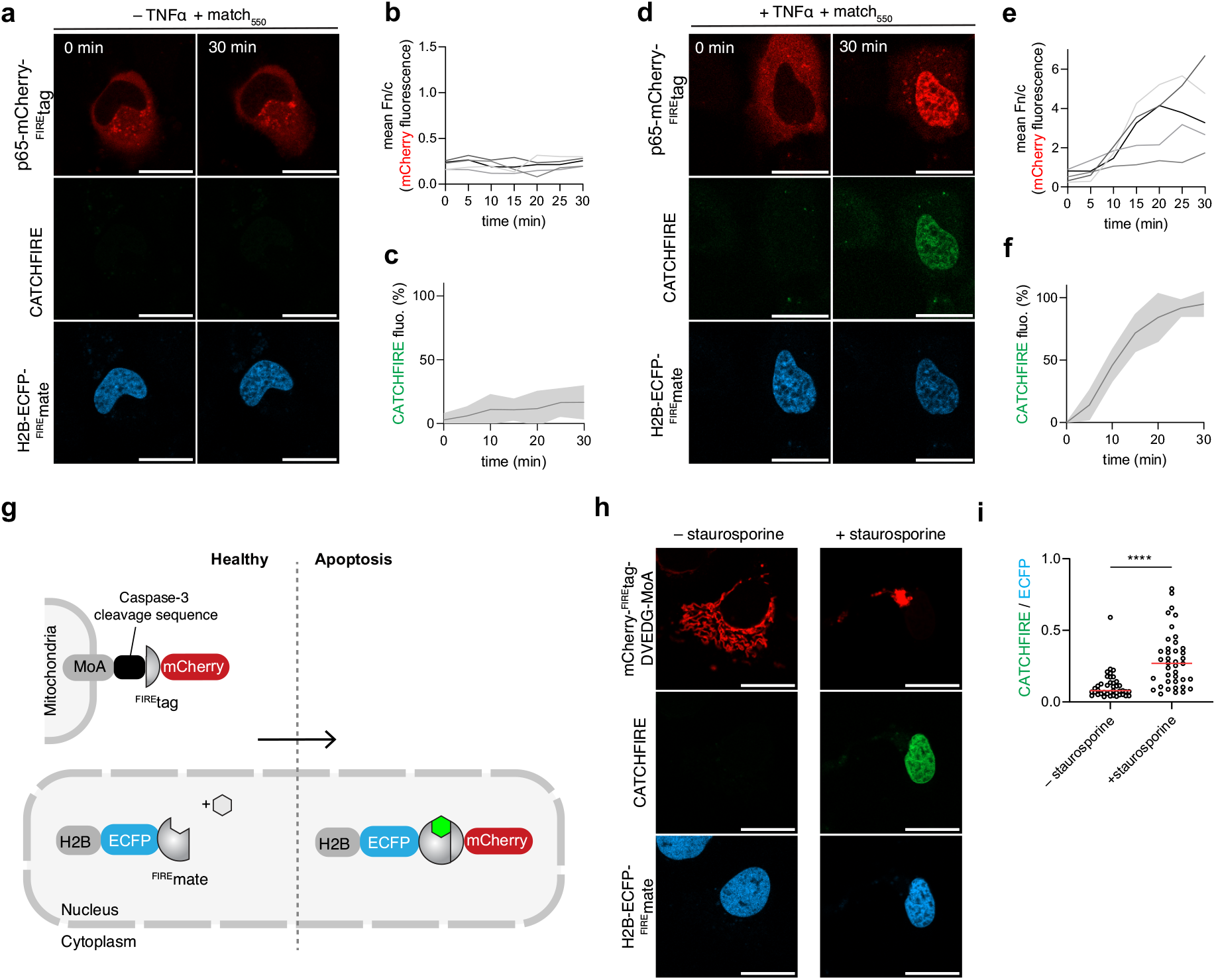
CATCHFIRE-based biosensors for studying cell signaling. **a-f** Signal transduction sensor. Hela cells expressing p65-mCherry-^FIRE^tag and H2B-ECFP-^FIRE^mate pretreated with 10 µM match_550_ were treated without (a-c) or with 10 ng/ml TNFα (d-f). **a**,**d** Representative micrographs of cells after 0 min and 30 min. Experiments were repeated three times with similar results. **b**,**e** Temporal evolution of the ratio nucleus-to-cytoplasm mCherry fluorescence of n = 5 cells (a-c) and n = 5 cells (d-f) from three independent experiments. **c**,**f** Temporal evolution of the CATCHFIRE signal. Data represent the mean ± SD of n = 5 cells (**a-c**) and n = 5 cells (**d-f**) from three independent experiments. **g** Schematic of the apoptosis sensor. **h**,**i** Hela cells expressing mCherry-^FIRE^tag-DVEDG-MoA and H2B-ECFP-^FIRE^mate pretreated with match_550_ were treated without or with 2μM staurosporine for 3.5 h. **h** Representative micrographs. Experiments were repeated three times with similar results. **i** Quantification of the apoptosis-induced CATCHFIRE. Data represent n = 41 cells (– staurosporine) and n = 40 cells (+ staurosporine) from two independent experiments. The median of CATCHFIRE/ECFP ratio is reported as a red line. An unpaired Student’s *t-*test allowed to compare the two distributions. **a**,**d**,**h** Red: ex/em 561/606-675 nm; green: ex/em 488/508-570 nm; cyan: ex/em 445/455-499 nm. Scale bars are 20 µm.

To explore the generality of this approach, we designed a sensor of apoptosis by relying on apoptosis-mediated nuclear translocation. We fused mCherry- ^FIRE^tag at the N-terminus of the mitochondrial monoamine oxidase (MoA) protein using a DEVDG linker and expressed it together with H2B-ECFP-^FIRE^mate as nuclear anchor. DEVDG sequence can be cleaved by caspase-3 during cell apoptosis, releasing thus mCherry- ^FIRE^tag. Prolonged treatment of cells with staurosporine resulted in the release of mCherry- ^FIRE^tag by action of the caspase-3 and its translocation to the nucleus where it interacted with H2B-ECFP-^FIRE^mate inducing a nuclear green fluorescence (**Fig. 6g-i**). The principle of such CATCHFIRE-based fluorescent sensors is general and could be applied to design optical sensors for various other cellular activities. As CATCHFIRE-based sensing does not require the use of microscopes, techniques such as flow cytometry or other fluorometric techniques could also be used for the detection.

## DISCUSSION

CATCHFIRE uses fluorogenic inducers of proximity to conditionally induce and visualize the proximity of two proteins fused to a small peptide tag (^FIRE^tag) and a small protein domain (^FIRE^mate). We show here that CATCHFIRE allows to develop both original inducers and sensors usable in living cells. Compared to traditional CIP assays, CATCHFIRE has several advantages. First, it allows both to induce and visualize the recruitment process without the need for additional reporters. Fluorogenic inducers of proximity with various spectral properties exist allowing to adjust the color to a given experimental set-up. Second, CATCHFIRE is the CIP tool with the smallest dimerizing domains (11 amino acids for ^FIRE^tag and 114 amino acids for ^FIRE^mate), which should reduce the risk of dysfunctional fusions. The small size of ^FIRE^tag may also enable fast genome editing for endogenous tagging. Third, the full reversibility of the fluorogenic recruitment upon washout of the fluorogenic inducer of proximity enables to control protein proximity with an unprecedent temporal resolution. Here again, simultaneous visualization of protein release is achieved without additional tagging.

In this study, we demonstrated the use of CATCHFIRE to control and track with high spatiotemporal resolution protein rerouting, nucleocytoplasmic protein trafficking, secretory protein trafficking, organelle transport and key cellular mechanisms such as mitophagy. Rerouting protein to a given organelle or cell compartment should allow in the future rapid tuning of cellular functions through reversible protein knock-sideway^23^.

CATCHFIRE could also be used to induce recruitment of proteins or protein domains and activate cellular functions. The fluorogenic nature of CATCHFIRE opens also unprecedented possibilities for the design of sensors with built-in multi-color fluorescence activation. As the efficiency of CATCHFIRE depends on the ability of ^FIRE^tag and ^FIRE^mate to meet each other, we could design sensors that switch on upon activation of signaling pathways or cellular mechanisms involving protein localization changes. We anticipate that CATCHFIRE will enable the development of novel sensors with built-in fluorescence activation that will allow to study a large diversity of cellular processes. Such set-up will be particularly well-suited for cell imaging but because of the fluorogenic nature of CATCHFIRE, assays independent from microscopy analysis (e.g. fluorometry, cytometry) can also be developed, opening great prospect for larger scale screenings.

## METHODS

### Data availability

All data are available in the article and supplementary information. Source data of all data presented in graphs within the figures are provided with this paper. The plasmids used in this study will be available from Addgene.

The presented research complies with all relevant ethical regulations.

### General

Synthetic oligonucleotides used for cloning were purchased from Integrated DNA technology. PCR reactions were performed with Q5 polymerase (New England Biolabs) in the buffer provided. PCR products were purified using QIAquick PCR purification kit (Qiagen). DNAse I, T4 ligase, Fusion polymerase, Taq ligase and Taq exonuclease were purchased from New England Biolabs and used with accompanying buffers and according to manufacturer protocols. Isothermal assemblies (Gibson assembly) were performed using homemade mix prepared according to previously described protocols^24^. Small-scale isolation of plasmid DNA was done using QIAprep miniprep kit (Qiagen) from 2 mL overnight bacterial culture supplemented with appropriate antibiotics. Large-scale isolation of plasmid DNA was done using the QIAprep maxiprep kit (Qiagen) from 150 mL of overnight bacterial culture supplemented with appropriate antibiotics. All plasmid sequences were confirmed by Sanger sequencing with appropriate sequencing primers (GATC Biotech). All the plasmids used in this study are listed in **Supplementary Table S1**, as well as DNA sequences. The synthesis of match_540_ (a.k.a. HMBR), match_550_ (a.k.a. HBR-2,5DM), match_600_ (a.k.a HBR-3,5DOM), match_dark_ (a.k.a. HBIR-3M) were previously reported^13,14,25^.

### Cloning

The plasmids used in this study have been generated using isothermal Gibson assembly or restriction enzymes and allow mammalian expression of relative insert under the control of the CMV promoter. The plasmid pAG1141 allowing the expression of FKBP-caged^FIRE^tag-IRES-IRFP670 was constructed by replacing the sequence coding for CFAST10 by the ^FIRE^tag sequence and the sequence of mCherry by the sequence of IRF670 in the vector pAG496^11^ coding for FKBP-CFAST10-IRES-mCherry. The plasmid pAG1151 allowing the expression of FKBP- ^FIRE^tag -IRES-IRFP670 was constructed by removing the sequence coding for PYP_2-114_ from the vector pAG1141. The plasmid pAG1142 allowing the expression of FRB-^FIRE^mate-IRES-mTurquoise2 was constructed by replacing the sequence coding for FAST_1-114_ by ^FIRE^mate sequence, in the previously described vector pAG64^11^. The plasmid pAG1159 allowing the expression of TOM20-ECFP-^FIRE^mate was constructed by replacing the sequence coding for FRB by ^FIRE^mate sequence in the Addgene vector #171461 Tom20-CR coding for TOM20-ECFP-FRB^25^. Tom20-CR was a gift from Takafumi Miyamoto (Addgene plasmid # 171461 ; http://n2t.net/addgene:171461 ; RRID:Addgene_171461). The plasmid pAG1160 allowing the expression of mCherry-^FIRE^tag was constructed by replacing the sequence coding for PYP by ^FIRE^tag sequence in the previously described vector pAG97^14^ coding for mCherry-PYP. The plasmid pAG1209 allowing the expression of ^FIRE^tag-mCherry was constructed by adding the sequence coding for ^FIRE^tag in the vector pAG1189 (Gautier et al. unpublished) coding for mCherry. The plasmid pAG1241 allowing the expression of ^FIRE^tag-mCherry-^FIRE^tag was constructed by adding the sequence coding for ^FIRE^tag in the vector pAG1209 coding for ^FIRE^tag-mCherry. The plasmid pAG1188 allowing the expression of mCherry-caged^FIRE^tag was constructed by adding the sequence coding for PYP_2-114_ in the vector pAG1160. The plasmid pAG1344 allowing the expression of EGFP-^FIRE^tag was constructed by replacing the sequence coding for mCherry by EGFP sequence in the vector pAG1160. The plasmid pAG1169 for the expression of ^FIRE^mate-ECFP-Cb5 was constructed by replacing the sequence coding for FKBP1C by ^FIRE^mate sequence in the Addgene vector (#162438) coding for FKBP1C-ECFP-Cb5^26^. FKBP1C-CFP-Cb5 was a gift from Takanari Inoue (Addgene plasmid # 162438 ; http://n2t.net/addgene:162438 ; RRID:Addgene_162438). The plasmid pAG1171 for the expression of ^FIRE^mate-ECFP-Giantin was constructed by replacing the sequence coding for FRB-ECFP(W66A)-Giantin by ^FIRE^mate-ECFP sequence in the Addgene vector (#67903) coding for FRB-ECFP(W66A)-Giantin^15^. FRB-ECFP(W66A)-Giantin was a gift from Dorus Gadella (Addgene plasmid # 67903 ; http://n2t.net/addgene:67903 ; RRID:Addgene_67903). The plasmid pAG1186 for the expression of Lyn11-ECFP-^FIRE^mate was constructed by replacing the sequence coding for TOM20 by Lyn11 sequence in the vector pAG1159. The plasmid pAG1211 for the expression of FRB-ECFP-Giantin was constructed by replacing the sequence coding for ECFP(W66A) by ECFP sequence in the Addgene vector (#67903) coding for FRB-ECFP(W66A)-Giantin^15^. The plasmid pAG1212 for the expression of mCherry-FKBP-^FIRE^tag was constructed by inserting the sequence coding for ^FIRE^tag in the vector pAG1190 coding for mCherry-FKBP. The plasmid pAG1242 for the expression of NLS-mCherry- ^FIRE^tag was constructed by inserting the sequence coding for NLS in the vector pAG1160. The plasmid pAG1243 for the expression of NES-ECFP-^FIRE^mate was constructed by replacing the sequence coding for TOM20 by NES sequence in the vector pAG1159. The plasmid pAG1280 for the expression of NLS-mCherry- ^FIRE^tag-NES was constructed by inserting the sequence coding for NES in the vector pAG1242. The plasmid pAG1247 for the expression of H2B-ECFP- ^FIRE^mate was constructed by replacing the sequence coding for TOM20 by H2B sequence in the vector pAG1159. The plasmid pAG1244 for the expression of p65-mCherry- ^FIRE^tag was constructed by inserting the sequence coding for p65, amplified from the Addgene vector GFP-RelA (#23255)^27^, in the vector pAG1160. GFP-RelA was a gift from Warner Greene (Addgene plasmid # 23255; http://n2t.net/addgene:23255; RRID:Addgene_23255). The plasmid pAG1237 for the expression of ^FIRE^mate-ECFP-Fis1 was constructed by inserting the sequence coding for Fis1, amplified form the Addgene vector pHAGE-mt-mKeima-P2A-FRB-Fis1 (#135295)^28^, in the vector pAG1165. pHAGE-mt-mKeima-P2A-FRB-Fis1 was a gift from Richard Youle (Addgene plasmid # 135295; http://n2t.net/addgene:135295; RRID:Addgene_135295). The plasmid pAG1238 for the expression of PINK1-mCherry-^FIRE^tag was constructed by inserting the sequence coding for PINK1, amplified from the Addgene vector pEYFP-N1-Pink1 (#101874)^29^, in the vector pAG1160. pEYFP-N1-Pink1 was a gift from Richard Youle (Addgene plasmid # 101874; http://n2t.net/addgene:101874; RRID:Addgene_101874). The plasmid pAG1345 for the expression of mCherry-^FIRE^tag-DEVDG-MoA was constructed by inserting the sequence coding for DEVDG-MoA, amplified from the Addgene vector ECFP(W66A)-FRB-MoA (#67904)^15^, in the vector pAG1160. ECFP(W66A)-FRB-MoA was a gift from Dorus Gadella (Addgene plasmid # 67904; http://n2t.net/addgene:67904; RRID:Addgene_67904). The plasmid pFP5290 for the expression of ^FIRE^mate-KDEL-IRES-mApple-^FIRE^tag-GPI was constructed by insertion of ^FIRE^tag-GPI obtained from gene synthesis (Integrated DNA technology) into a vector for the expression of Streptavidin-KDEL-IRES-SBP-mApple-GPI (built from the plasmid pFP1774^17^, Addgene plasmid # 65294; http://n2t.net/addgene:65294; RRID:Addgene_65294; see also Addgene plasmid #166904, http://n2t.net/addgene:166904), and then insertion of ^FIRE^mate obtained from gene synthesis (Integrated DNA technology) instead of Streptavidin. The plasmid pFP5299 for the expression of ^FIRE^mate-KDEL-IRES-ManII-mApple-^FIRE^tag was constructed by addition of ^FIRE^tag into a vector for the expression of Streptavidin-KDEL-IRES-ManII-SBP-mApple (constructed from pFP1708^17^, Addgene plasmid # 65252; http://n2t.net/addgene:65252; RRID:Addgene_65252) and then insertion of ManII-SBP-mApple-^FIRE^tag into pFP5290 instead of mApple-^FIRE^tag-GPI. The plasmid pFP5370 for the expression of ^FIRE^mate-KDEL-IRES-TNF-mCherry-^FIRE^tag was constructed by insertion of ^FIRE^tag in pFP1752^17^, (Addgene plasmid # 65279; http://n2t.net/addgene:65279; RRID:Addgene_65279) and then insertion of TNF-mCherry-FIREtag into pFP5290 instead of mApple-^FIRE^tag-GPI. The plasmid pFP5369 for the expression of KIF17MD-flag-^FIRE^mate was constructed by insertion of ^FIRE^mate obtained from gene synthesis (Integrated DNA technology) into a vector for the expression of KIF17MD-flag-FRB (constructed from a vector given by Lukas Kapitein^20^) instead of FRB. The plasmid pFP5380 for the expression of LAMP1-mCherry-^FIRE^tag was constructed by insertion of mCherry-^FIRE^tag from pFP5370 in a plasmid for the expression of LAMP1-SBP-EGFP.

### Cell culture

HeLa cells (ATCC CRM-CCL2) were cultured in Minimal Essential Media (MEM) supplemented with phenol red, Glutamax I, 1 mM of sodium pyruvate, 1% (vol/vol) of non-essential amino-acids and 10% (vol/vol) fetal calf serum (FCS), at 37 °C in a 5% CO_2_ atmosphere. HEK 293T (ATCC CRL-3216) cells were cultured in Dulbecco’s Modified Eagle Medium (DMEM) supplemented with phenol red and 10% (vol/vol) fetal calf serum at 37 °C in a 5% CO_2_ atmosphere. U2OS cells (ATCC HTB-96) were cultured in Dulbecco’s modified Eagle’s medium (DMEM) supplemented with phenol red and 10% (vol/vol) fetal calf serum and 1% (vol/vol) penicillin–streptomycin at 37°C in a 5% CO_2_ atmosphere. For imaging, cells were seeded in µDish IBIDI (Biovalley) coated with poly-L-lysine. The ratios of plasmids used for each experiment are given in **Table S2**. Cells were transiently transfected using Genejuice (Merck) according to the manufacturer’s protocols for 24 prior to imaging. Cells were washed with DPBS (Dulbecco’s Phosphate-Buffered Saline), and treated with DMEM media (without serum and phenol red) supplemented with the compounds at the indicated concentration. For the p65 nuclear translocation experiments, cells were serum-starved for 4 hours before activation by TNFα. For experiments on reversibility, secretory protein trafficking and organelle positioning, HeLa cells were cultured in DMEM supplemented with 10% FCS, 1 mM sodium pyruvate and 100 µg/mL penicillin and streptomycin at 37°C in 5% CO_2_ atmosphere. HeLa cells were transfected using calcium phosphate as previously described^30^. For imaging, cells were seeded onto 25 mm-diameter glass coverslips. The day after transfection, coverslips were transferred into L-shape tubing Chamlide (Live Cell Instrument) filled with carbonate independent Leibovitz’s medium supplemented with 10% FCS, and the compounds at the indicated concentration. Washouts were done by addition into the Chamlide of pre-warmed PBS (Phosphate-Buffered Saline) 2 mL at a time for a total volume of 15 mL, and then addition of Leibovitz’s medium.

### Flow cytometry analysis

Flow cytometry in HEK 293T cells was performed on a MACSQuant® analyzer equipped with 405 nm, 488 nm and 635 nm lasers and seven channels. To prepare samples, cells were first grown in cell culture flasks, then transiently co-transfected 24 hours after seeding using Genejuice (Merck) according to the manufacturer’s protocol for 24 hours. After 24 hours, cells were centrifuged in PBS with BSA (1 mg/ml) and resuspend in PBS-BSA supplemented with the appropriate amounts of compounds. For each experiment, 20,000 cells positively expressing mTurquoise2 (Ex 434 / Em 450 ± 25) and iRFP670 (Ex 638 nm / Em 660 ± 10 nm) were analyzed with the following parameters: Ex 488nm, Em 525 ± 20 nm. Data were analyzed using FlowJo v10.7.1.

### Fluorescence microscopy

The confocal micrographs were acquired on a Zeiss LSM 980 Laser Scanning Microscope equipped with a plan apochromat 63× /1.4 NA oil immersion objective. ZEN software was used to collect the data. Confocal spinning disk images were acquired on a Nikon Inverted Eclipse Ti-E (Nikon) microscope equipped with a Spinning disk CSU-X1 (Yokogawa), an iXon EMCCD camera (Andor) integrated in Metamorph software by Gataca Systems, using an 60× CFI plan apochromat VC 1.4 NA oil immersion objective (Nikon). The images were analyzed using Icy (2.4.0.0), Fiji (Image J) and NIS-Elements (Nikon). Images from figures 3 and 4 were enhanced by Denoise.ai on NIS-Elements Ver.5.42 and 2D-deconvolution on NIS-Elements Ver.5.42. To track fluorescence signal in a specific organelle, the organelle contour was determined by masking the signal in the ECFP channel with the plug-in *HK-means* with the following parameters: intensity class equals 100, min object size (px) equals 500, max object size (px) equals 1500-3000. The signal intensity of the ROI was tracked over the time for each channel by using plug-in *Active Contours*. The background signal was subtracted. Data were processed using GraphPad Prism v9.3.0.

## ACKNOWLEDGMENTS

We thank the imaging facility of the Institut de Biologie Paris Seine of Sorbonne University as well as the PICT imaging facility of the Institut Curie. We acknowledge Takafumi Miyamoto for the plasmid Tom20-CR (Addgene plasmid # 171461) Takanari Inoue for the plasmid FKBP1C-CFP-Cb5 (Addgene plasmid #162438), Dorus Gadella for the plasmids FRB-ECFP(W66A)-Giantin (Addgene plasmid # 67903) and ECFP(W66A)-FRB-MoA (Addgene plasmid # 67904), Warner Greene for the plasmid GFP-RelA (Addgene plasmid # 23255), Richard Youle for the plasmids pHAGE-mt-mKeima-P2A-FRB-Fis1 (Addgene plasmid # 135295) and pEYFP-N1-Pink1 (Addgene plasmid # 101874). The work in AG lab has been supported by the European Research Council (ERC-2016-CoG-724705 FLUOSWITCH), the Agence Nationale de la Recherche (ANR-19-CE13-0026 ADOBE) and the Institut Universitaire de France. The work in FP lab has been supported by the Fondation pour la Recherche Médicale (EQU201903007925), and the Agence Nationale de la Recherche (ANR-19-CE13-0006-03; ANR-20-CE14-0017-02; ANR-19-CE13-0002-03; ANR-11-LABX-0038), and has also received support under the program « Investissements d’Avenir » launched by the French Government and implemented by ANR with the references CelTisPhyBio (11-LBX-0038) and ANR-10-IDEX-0001-02 PSL.

## AUTHOR CONTRIBUTIONS

S. B., Z.V.C., O.J., G.B., F. P. and A.G. designed the experiments. S.B., Z.V.C., O.J. and G.B. performed the experiments. S. B., Z.V.C., O.J., G.B., F. P. and A.G. analyzed the experiments. S. B., F. P. and A.G. wrote the paper with the help of all the authors.

