## Supplementary information for "A fluorogenic chemically induced dimerization technology for controlling, imaging and sensing protein proximity"

**This PDF file includes:**

- Legends of Movies S1-S23
- Fig. S1 to S10
- Tables S1 to S2
- Supplementary references

#### Legends of Supplementary Movies

**Movie S1. Fluorogenic rerouting of cytosolic proteins to mitochondria with match$_{550}$.** HeLa cells co-expressing mCherry-$^{FIRE}$tag together with Tom20-ECFP-$^{FIRE}$mate were treated with 10 μM match$_{550}$, and imaged by time-lapse confocal microscopy (see also **Fig. 1c,d**). Experiments were repeated three times with similar results. Red (mCherry): ex/em 561/606-675 nm; Green (CATCHFIRE): ex/em 488/508-570 nm; Cyan (ECFP): ex/em 445/455-499 nm; Scale bars are 20 μm.

**Movie S2. Kinetics of CATCHFIRE**. HeLa cells co-expressing mCherry-$^{FIRE}$tag together with Tom20-ECFP-$^{FIRE}$mate were treated with 10 μM match$_{550}$, and imaged by fast time-lapse confocal microscopy (see also **Fig. S3a,b**). Experiments were repeated three times with similar results. Green (CATCHFIRE): ex/em 488/508-570 nm; Scale bars are 20 μm.

**Movie S3. Stability of CATCHFIRE**. U2OS cells co-expressing mCherry-$^{FIRE}$tag and Tom20-ECFP-$^{FIRE}$mate were treated with match$_{550}$ and image by time-lapse confocal microscopy at 1 image every 2 minutes (see also **Fig. S3c,d**). Experiment was repeated three times with similar results. Red (mCherry): ex/em 561/606-675 nm; Green (CATCHFIRE): ex/em 488/508-570 nm; Cyan (ECFP): ex/em 445/455-499 nm; Scale bars are 20 μm.

**Movie S4. Fluorogenic rerouting of cytosolic proteins to mitochondria with match$_{550}$.** HeLa cells co-expressing $^{FIRE}$tag-mCherry together with Tom20-ECFP-$^{FIRE}$mate were treated with 10 μM match$_{550}$, and imaged by time-lapse confocal microscopy (see also **Fig. S4a,b**). Experiments were repeated three times with similar results. Red (mCherry): ex/em 561/606-675 nm; Green (CATCHFIRE): ex/em 488/508-570 nm; Cyan (ECFP): ex/em 445/455-499 nm; Scale bars are 20 μm.

**Movie S5. Fluorogenic rerouting of cytosolic proteins to mitochondria with match$_{550}$.** HeLa cells co-expressing mCherry-$^{FIRE}$tag-mCherry together with Tom20-ECFP-$^{FIRE}$mate were treated with 10 μM match$_{550}$, and imaged by time-lapse confocal microscopy (see also **Fig. S4c,d**). Experiments were repeated three times with similar results. Red (mCherry): ex/em 561/606-675 nm; Green (CATCHFIRE): ex/em 488/508-570 nm; Cyan (ECFP): ex/em 445/455-499 nm; Scale bars are 20 μm.

**Movie S6. Fluorogenic rerouting of cytosolic proteins to mitochondria with match$_{540}$.** HeLa cells co-expressing mCherry-$^{FIRE}$tag together with Tom20-ECFP-$^{FIRE}$mate were treated with 10 μM match$_{540}$, and imaged by time-lapse confocal microscopy (see also **Fig. 1e,f**). Experiments were repeated three times with similar results. Red (mCherry): ex/em 561/606-675 nm; Green (CATCHFIRE): ex/em 488/508-570 nm; Cyan (ECFP): ex/em 445/455-499 nm; Scale bars are 20 μm.

**Movie S7. Fluorogenic rerouting of cytosolic proteins to mitochondria with match$_{600}$.** HeLa cells co-expressing EGFP-$^{FIRE}$tag together with Tom20-ECFP-$^{FIRE}$mate were treated with 10 μM match$_{600}$, and imaged by time-lapse confocal microscopy (see also **Fig. 1g,h**). Experiments were repeated three times with similar results. Red (mCherry): ex/em 561/606-675 nm; Green (CATCHFIRE): ex/em 488/508-570 nm; Cyan (ECFP): ex/em 445/455-499 nm; Scale bars are 20 μm.

**Movie S8. Fluorogenic rerouting of cytosolic proteins to mitochondria with match$_{\mathbf{dark}}$.** HeLa cells co-expressing mCherry-$^{FIRE}$tag together with Tom20-ECFP-$^{FIRE}$mate were treated with 10 μM match$_{dark}$, and imaged by time-lapse confocal microscopy (see also **Fig. 1i,j**). Experiments were repeated three times with similar results. Red (mCherry): ex/em 561/606-675 nm; Green (CATCHFIRE): ex/em 488/508-570 nm; Cyan (ECFP): ex/em 445/455-499 nm; Scale bars are 20 μm.

**Movie S9. Fluorogenic rerouting of cytosolic proteins to the Golgi apparatus with match$_{\mathbf{550}}$.** HeLa cells co-expressing mCherry-$^{FIRE}$tag together with $^{FIRE}$mate-ECFP-Giantin were treated with 10 μM match$_{550}$, and imaged by time-lapse confocal microscopy (see also **Fig. 1k,l**). Experiments were repeated three times with similar results. Red (mCherry): ex/em 561/606-675 nm; Green (CATCHFIRE): ex/em 488/508-570 nm; Cyan (ECFP): ex/em 445/455-499 nm; Scale bars are 20 μm.

**Movie S10. Fluorogenic rerouting of cytosolic proteins to the endoplasmic reticulum with match$_{\mathbf{550}}$.** HeLa cells co-expressing mCherry-$^{FIRE}$tag together with $^{FIRE}$mate-ECFP-Cb5 were treated with 10 μM match$_{550}$, and imaged by time-lapse confocal microscopy (see also **Fig. 1m,n**). Experiments were repeated three times with similar results. Red (mCherry): ex/em 561/606-675 nm; Green (CATCHFIRE): ex/em 488/508-570 nm; Cyan (ECFP): ex/em 445/455-499 nm; Scale bars are 20 μm.

**Movie S11. Fluorogenic rerouting of cytosolic proteins to the plasma membrane with match$_{\mathbf{550}}$.** HeLa cells co-expressing mCherry-$^{FIRE}$tag together with Lyn11-$^{FIRE}$mate-ECFP were treated with 10 μM match$_{550}$, and imaged by time-lapse confocal microscopy (see also **Fig. 1o,p**). Experiments were repeated three times with similar results. Red (mCherry): ex/em 561/606-675 nm; Green (CATCHFIRE): ex/em 488/508-570 nm; Cyan (ECFP): ex/em 445/455-499 nm; Scale bars are 20 μm.

**Movie S12. Fluorogenic rerouting of cytosolic proteins to the Golgi apparatus with match$_{\mathbf{540}}$.** HeLa cells co-expressing mCherry-$^{FIRE}$tag together with $^{FIRE}$mate-ECFP-Giantin were treated with 10 μM match$_{540}$, and imaged by time-lapse confocal microscopy (see also **Fig. S5a,b**). Experiments were repeated three times with similar results. Red (mCherry): ex/em 561/606-675 nm; Green (CATCHFIRE): ex/em 488/508-570 nm; Cyan (ECFP): ex/em 445/455-499 nm; Scale bars are 20 μm.

**Movie S13. Fluorogenic rerouting of cytosolic proteins to the endoplasmic reticulum with match$_{\mathbf{540}}$.** HeLa cells co-expressing mCherry-$^{FIRE}$tag together with $^{FIRE}$mate-ECFP-Cb5 were treated with 10 μM match$_{540}$, and imaged by time-lapse confocal microscopy (see also **Fig. S5c,d**). Experiments were repeated three times with similar results. Red (mCherry): ex/em 561/606-675 nm; Green (CATCHFIRE): ex/em 488/508-570 nm; Cyan (ECFP): ex/em 445/455-499 nm; Scale bars are 20 μm.

**Movie S14. Fluorogenic rerouting of cytosolic proteins to the plasma membrane with match$_{540}$.** HeLa cells co-expressing mCherry-$^{FIRE}$tag together with Lyn11-$^{FIRE}$mate-ECFP were treated with 10 μM match$_{540}$, and imaged by time-lapse confocal microscopy (see also **Fig. S5e,f**). Experiments were repeated three times with similar results. Red (mCherry): ex/em 561/606-675 nm; Green (CATCHFIRE): ex/em 488/508-570 nm; Cyan (ECFP): ex/em 445/455-499 nm; Scale bars are 20 μm.

**Movie S15. Fluorogenic recruitment of proteins is reversible through washout.** HeLa cells co-expressing mCherry-$^{FIRE}$tag and Giantin-ECFP-$^{FIRE}$mate were treated with 10 μM match$_{550}$, washed and then treated with 10 μM match$_{dark}$. Cells were imaged by time-lapse spinning-disk microscopy (See also **Fig. 1q**). Experiments were repeated three times with similar results. Red (mCherry): ex/em 561/604-664 nm; Green (CATCHFIRE): ex/em 488/525-545 nm; Scale bars are 10 μm.

**Movie S16. Fluorogenic recruitment of proteins with caged$^{FIRE}$tag.** HeLa cells co-expressing mCherry-caged$^{FIRE}$tag and $^{FIRE}$mate-ECFP-Giantin were treated with 10 μM match$_{550}$, and imaged by time-lapse confocal microscopy (see also **Fig. S8c,d**). Experiments were repeated three times with similar results. Red: ex/em 561/606-675 nm; green: ex/em 488/508-570 nm; cyan: ex/em 445/455-499 nm. Scale bars are 20 μm.

**Movie S17. Fluorogenic induction of nuclear export.** HeLa cells co-expressing NLS-mCherry-$^{FIRE}$tag and NES-ECFP- $^{FIRE}$mate were treated with 10 μM match$_{550}$ and imaged by time-lapse confocal microscopy (see also **Fig. 2a-d**). Experiments were repeated three times with similar results. Red: ex/em 561/606-675 nm; green: ex/em 488/508-570 nm; cyan: ex/em 445/455-499 nm. Scale bars are 20 μm.

**Movie S18. Fluorogenic induction of nuclear import.** HeLa cells co-expressing NLS-mCherry-$^{FIRE}$tag-NES and H2B-ECFP- $^{FIRE}$mate were treated with 10 μM match$_{550}$ and imaged by time-lapse confocal microscopy (see also **Fig. 2e-h**). Experiments were repeated three times with similar results. Red: ex/em 561/606-675 nm; green: ex/em 488/508-570 nm; cyan: ex/em 445/455-499 nm. Scale bars are 20 μm.

**Movie S19. Control of secretory protein trafficking.** HeLa cells co-expressing $^{FIRE}$mate-KDEL and TNF-mCherry-$^{FIRE}$tag were treated with match$_{550}$ for 24 h and imaged by spinning-disk microscopy after washout of match$_{550}$ (see also **Fig. 3b**). Experiments were repeated three times with similar results. Red (mCherry): ex/em 561/604-664 nm; Green (CATCHFIRE): ex/em 488/525-545 nm; Scale bars are 10 μm. Images were enhanced by Denoise.ai on NIS-Elements Ver.5.42 and by 2D-deconvolution on NIS-Elements Ver.5.42.

**Movie S20. Control of secretory protein trafficking.** HeLa cells co-expressing $^{FIRE}$mate-KDEL and mApple-$^{FIRE}$tag-GPI were treated with match$_{550}$ for 24 h and imaged by spinning-disk microscopy after washout of match$_{550}$ (see also **Fig. 3c**). Experiments were repeated three times with similar results. Red (mApple): ex/em 561/604-664 nm; Green (CATCHFIRE): ex/em 488/525-545 nm; Scale bars are 10 μm. Images were enhanced by Denoise.ai on NIS-Elements Ver.5.42 and by 2D-deconvolution on NIS-Elements Ver.5.42.

**Movie S21. Control of secretory protein trafficking.** HeLa cells co-expressing $^{FIRE}$mate-KDEL and ManII-mApple-$^{FIRE}$tag were treated with match$_{550}$ for 24 h and imaged by spinning-disk microscopy after washout of match$_{550}$ (see also **Fig. 3d**). Match$_{550}$ was re-added after 50 min. Experiments were done in 25 μg/mL cycloheximide. Experiments were repeated four times with similar results. Red (mApple): ex/em 561/604-664 nm; Green (CATCHFIRE): ex/em 488/525-545 nm; Scale bars are 10 μm. Images were enhanced by Denoise.ai on NIS-Elements Ver.5.42 and by 2D-deconvolution on NIS-Elements Ver.5.42.

**Movie S22. Control and tracking of organelle positioning.** HeLa cells co-expressing LAMP1-mCherry-$^{FIRE}$tag and $^{FIRE}$mate-KIF17 were treated with match$_{550}$ and imaged by spinning-disk microscopy. After 60 min, match$_{550}$ was washed out and cells were imaged for 60 min without match$_{550}$, before re-addition of match$_{550}$ for 60 min (see also **Fig. 4**). Experiments were repeated four times with similar results. Red (mCherry): ex/em 561/604-664 nm; Green (CATCHFIRE): ex/em 488/525-545 nm; Scale bars are 10 μm. Images were enhanced by Denoise.ai on NIS-Elements Ver.5.42 and by 2D-deconvolution on NIS-Elements Ver.5.42.

**Movie S23. Fluorogenic rerouting of PINK1 to the outer mitochondrial membrane.** HeLa cells co-expressing ECFP-$^{FIRE}$mate-Fis1 and PINK1-mCherry-$^{FIRE}$tag were treated without or with 10 μM match$_{550}$ and imaged by time-lapse confocal microscopy (see also **Fig. S9**). Experiments were repeated three times with similar results. Red: ex/em 561/606-675 nm; green: ex/em 488/508-570 nm; cyan: ex/em 445/455-499 nm. Scale bars are 20 μm.

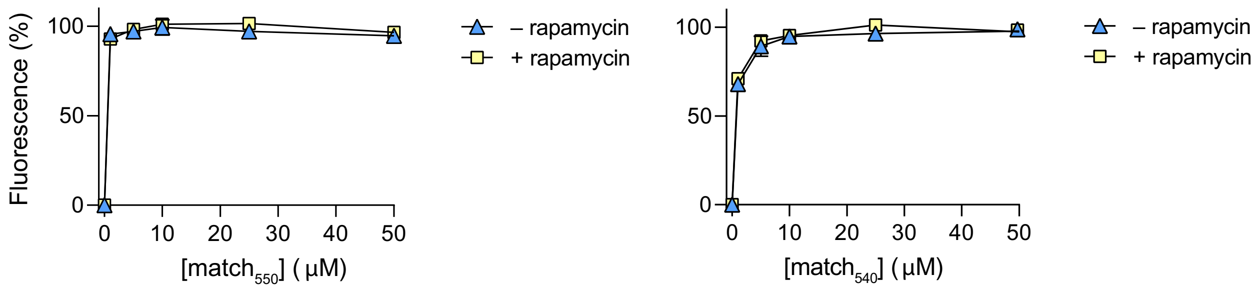


**Figure S1. Match-induced complementation of FIREmate (a.k.a. pFAST$_{1\text{-}114}$) and FIREtag (a.k.a. pFAST$_{115\text{-}125}$).** Normalized average fluorescence of about 20,000 HEK293T cells co-expressing the FK506-binding protein (FKBP) fused to FIREtag and the FKBP-rapamycin-binding domain of mammalian target of rapamycin (FRB) fused to FIREmate treated without or with 500 nM of rapamycin, and with 1, 5, 10, 25 or 50 µM of match$_{550}$ (a.k.a. HBR-2,5DM) and match$_{540}$ (a.k.a. HMBR). Individual cell fluorescence was analyzed by flow cytometry. Data represent the mean values ± SD of three independent experiments.

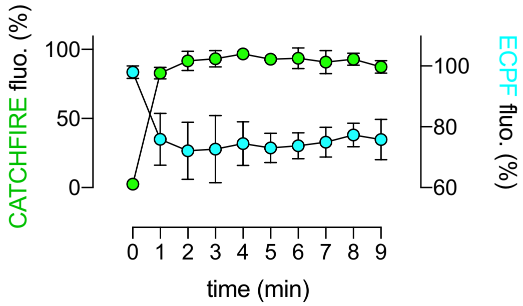


**Figure S2. FRET between ECFP and CATCHFIRE leads to a loss of ECFP fluorescence.** When using ECFP as an additional reporter of localization, we observed a decrease of the ECFP signal upon induction of CATCHFIRE, which we attributed to a loss of fluorescence by Förster resonance energy transfer between ECFP and the CATCHFIRE-induced ternary fluorescent assembly. As an example, the graph shows the temporal evolution of the ECFP fluorescence signal (right axis) upon CATCHFIRE induction (left axis) in HeLa cells co-expressing mCherry-$^{FIRE}$tag and Tom20-ECFP-$^{FIRE}$mate treated with 10 μM match$_{550}$ (data shown on **Fig. 1c,d**). Data represents the mean ± SD of n = 20 cells from three independent experiments.

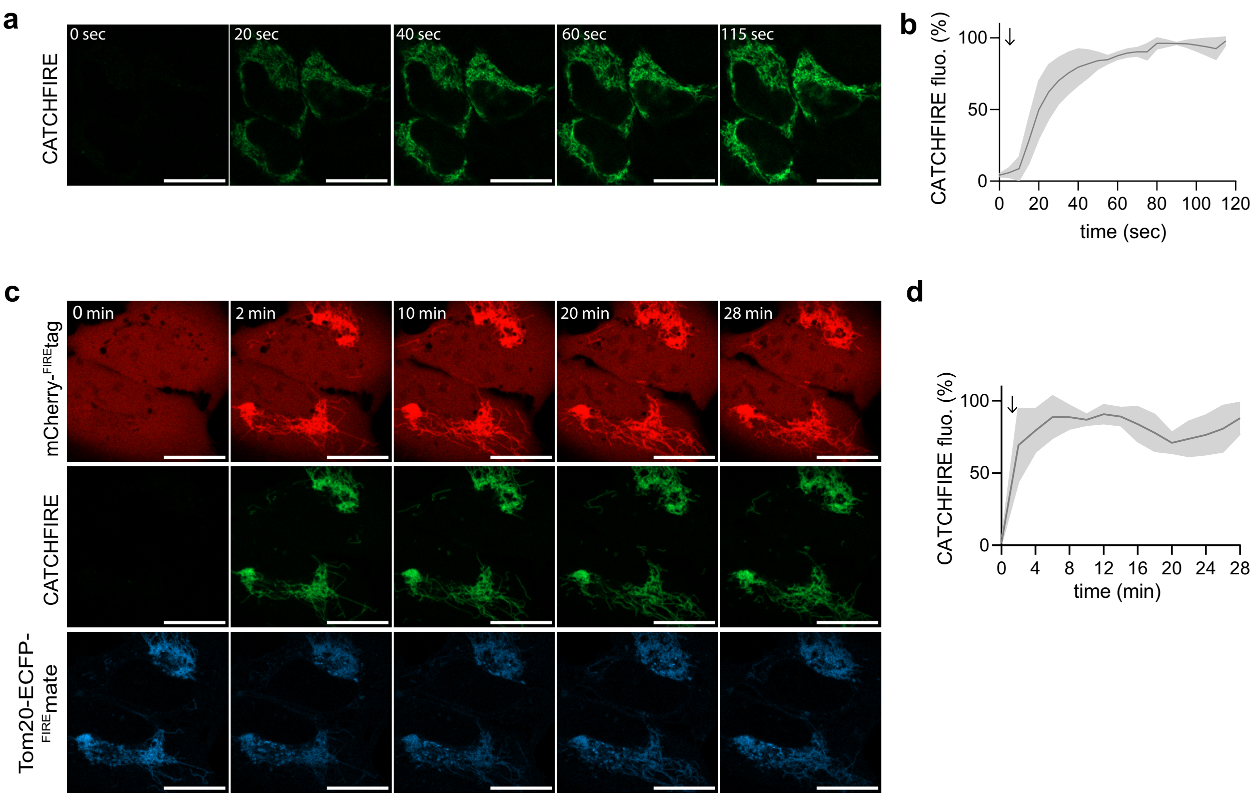


**Figure S3. Fluorogenic induced recruitment of cytoplasmic proteins to mitochondria. a,b** HeLa cells co-expressing mCherry-$^{FIRE}$tag and Tom20-ECFP-$^{FIRE}$mate were treated with 10 μM match$_{550}$ and imaged by time-lapse confocal microscopy at 1 image per 5 seconds. **a** Representative time-lapse of FIRE (ex/em 488/508-570 nm) (see also **Movie S2**). Experiment was repeated three times with similar results. Scale bars are 20 μm. **b** Temporal evolution of the FIRE signal. Data represents the mean values ± SD of 15 cells from three independent experiments. **c,d** U2OS cells co-expressing mCherry-$^{FIRE}$tag and Tom20-ECFP-$^{FIRE}$mate were treated with match$_{550}$ and image by time-lapse confocal microscopy at 1 image every 2 minutes. **c** Representative time-lapse (mCherry: ex/em 561/606-675 nm; FIRE: ex/em 488/508-570 nm; ECFP: ex/em 445/455-499 nm) (see also **Movie S3**). Experiment was repeated three times with similar results. Scale bars are 20 μm. **d** Temporal evolution of the CATCHFIRE signal. Data represents the mean values ± SD of 10 cells from three independent experiments.

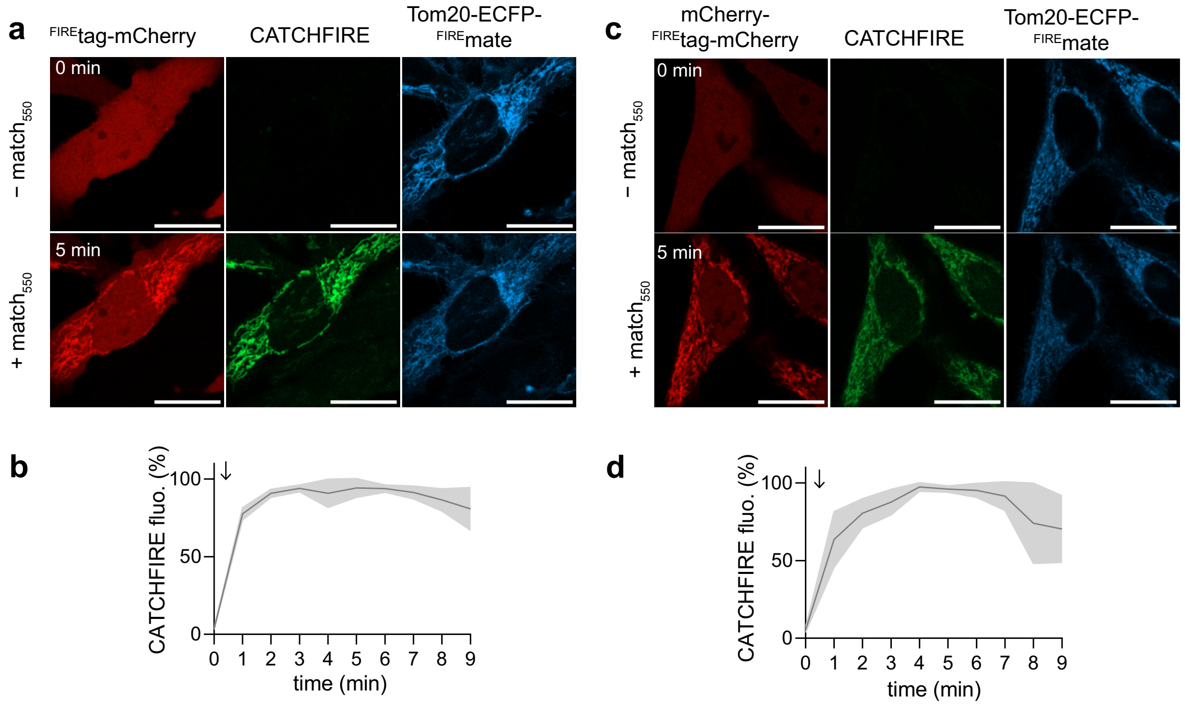


**Figure S4. $^{FIRE}$tag position does not influence CATCHFIRE efficiency.** HeLa cells co-expressing $^{FIRE}$tag-mCherry (a,b) or mCherry-$^{FIRE}$tag-mCherry (c,d) and Tom20-ECFP-$^{FIRE}$mate were treated with 10 μM $match_{550}$ and imaged by time-lapse confocal microscopy. **a,c** Representative confocal micrographs of cells before (0 min) and after (5 min) addition of $match_{550}$ (mCherry: ex/em 561/606-675 nm; CATCHFIRE: ex/em 488/508-570 nm; ECFP: ex/em 445/455-499 nm) (see also **Movies S4-S5**). Experiments were repeated three times with similar results. Scale bars are 20 μm. **b,d** Temporal evolution of the CATCHFIRE signal. Data represents the mean values ± SD of 17 cells (**a,b**) and 13 cells (**c,d**) from three independent experiments.

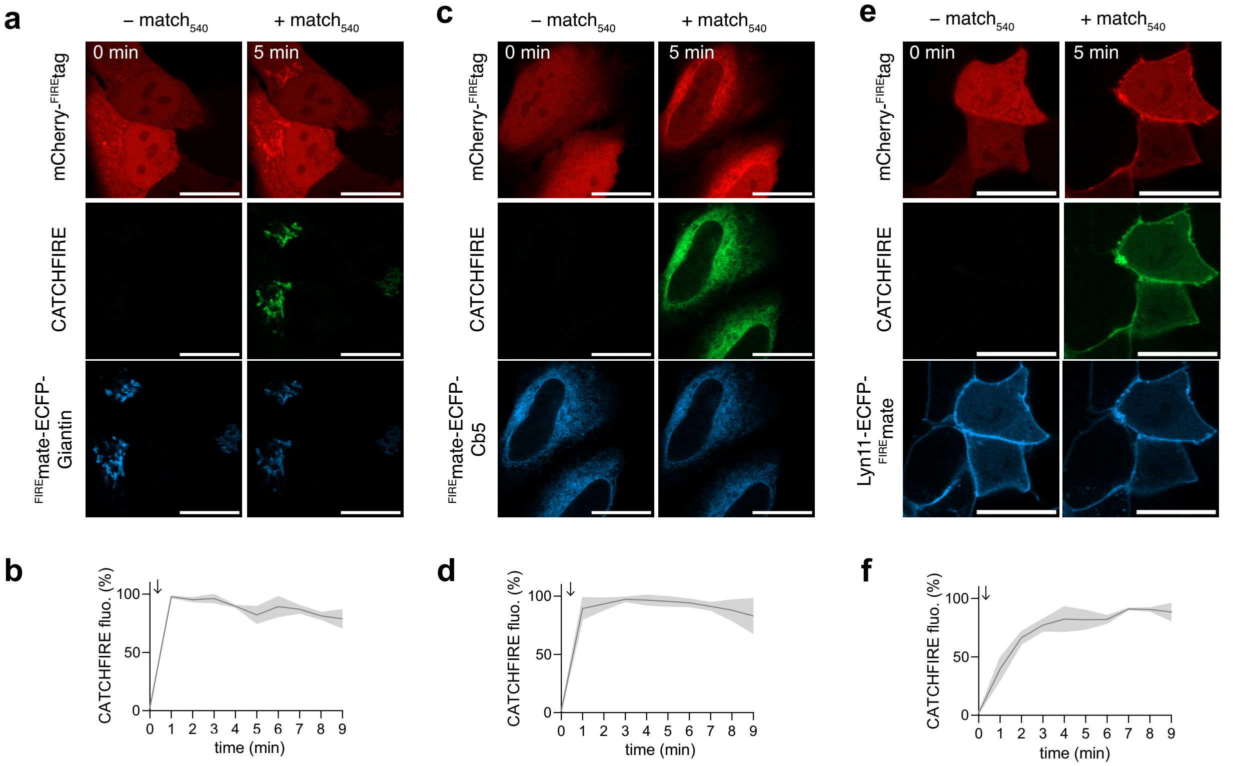


**Figure S5. CATCHFIRE to various organelles.** HeLa cells co-expressing mCherry-$^{FIRE}$tag and $^{FIRE}$mate-ECFP-Giantin (**a**,**b**), $^{FIRE}$mate-ECFP-Cb5 (**c**,**d**), Lyn11-$^{FIRE}$mate-ECFP (**e**,**f**) were treated with 10 μM $match_{540}$, and imaged by time-lapse confocal microscopy. **a,c,e** Representative confocal micrographs of cells before (0 min) and after (5 min) addition of $match_{540}$ (see **Movies S12-S14**). Experiments were repeated three times with similar results. **b,d,f** Temporal evolution of the CATCHFIRE signal. Data represents the mean values ± SD of 12 cells (**b**), 18 cells (**d**), 30 cells (**f**) from three independent experiments.

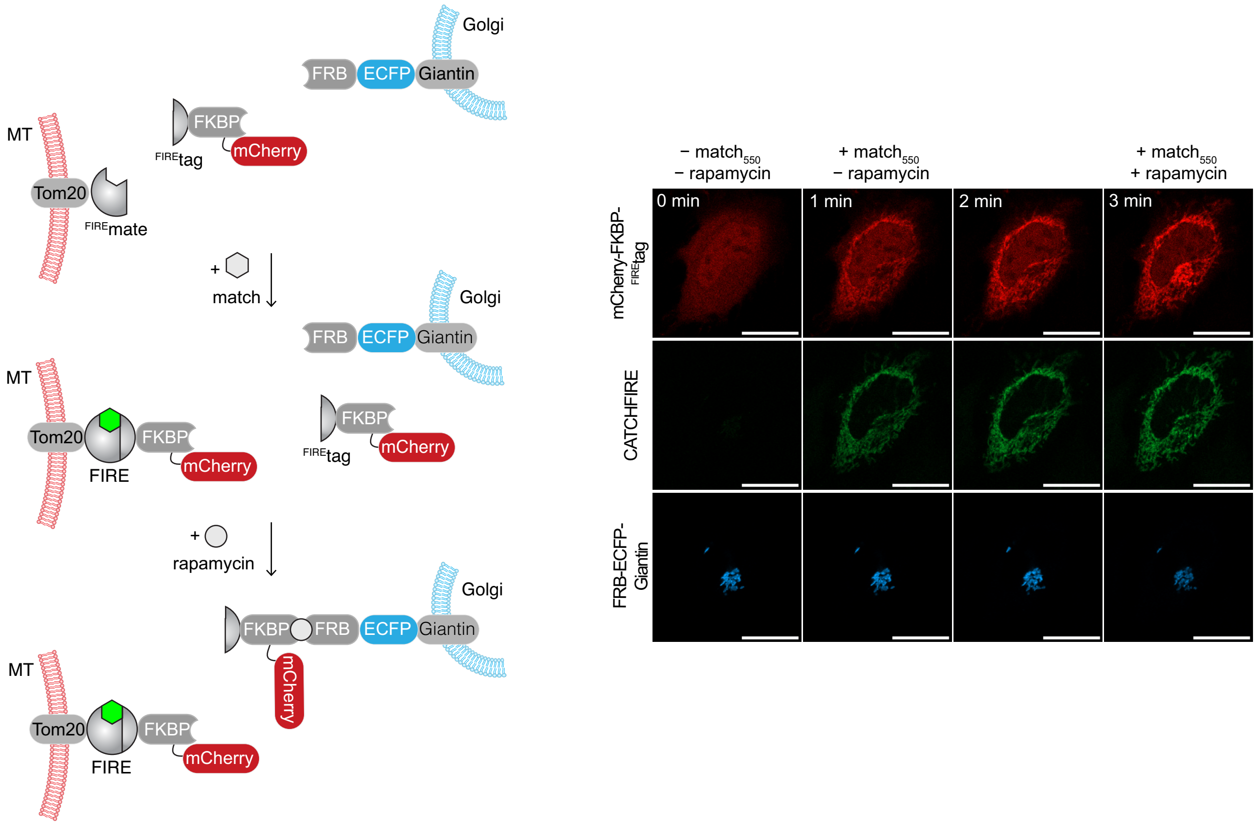


**Figure S6. Dual control with CATCHFIRE and FRB-rapamycin-FKBP.** HeLa cells co-expressing mCherry-FKBP-$^{FIRE}$tag, TOM20-$^{FIRE}$mate and FRB-ECFP-Giantin were treated with 10 μM match$_{550}$ and then, after 2 min, with 500 nM rapamycin, and imaged by time-lapse confocal microscopy. Representative confocal micrographs of cells before, after addition of match$_{550}$, and after addition of rapamycin. Red: ex/em 561/606-675 nm; green: ex/em 488/508-570 nm; cyan: ex/em 445/455-499 nm. Scale bars are 20 μm.

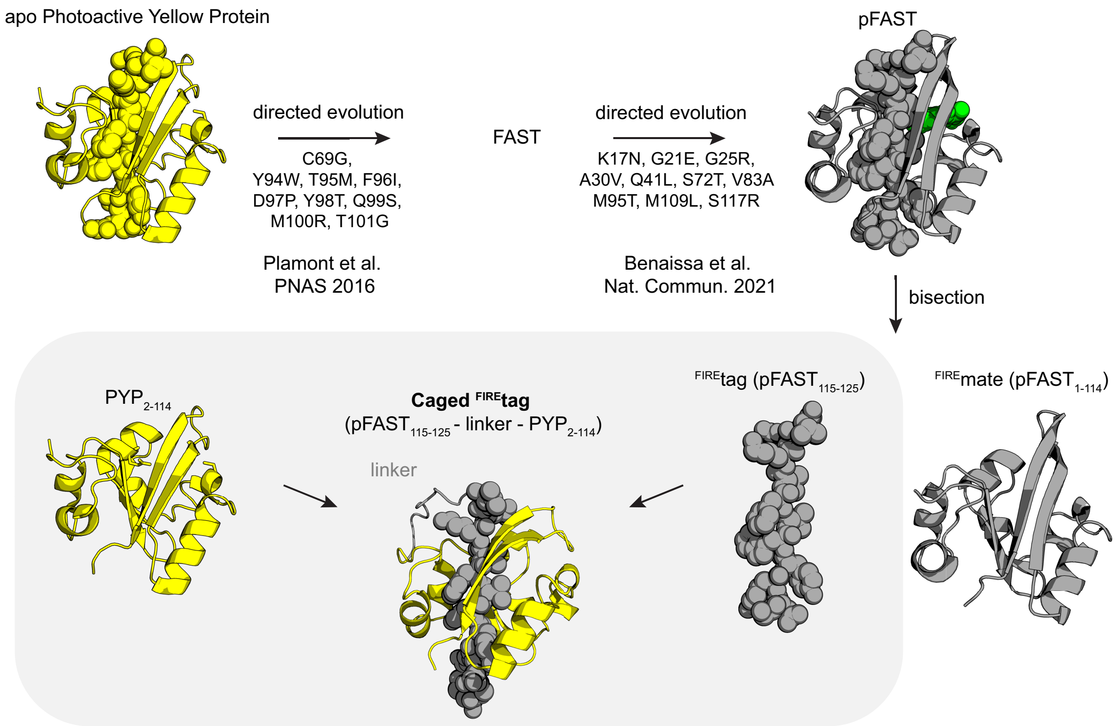


**Figure S7. Design of caged$^{FIRE}$tag.** Fusion of $^{FIRE}$tag at the N-terminal domain of the apo photoactive yellow protein ($PYP_{2\text{-}114}$) resulted in a protein (caged$^{FIRE}$tag) that folds like a circular permutation of PYP because of the intramolecular interaction between $^{FIRE}$tag and $PYP_{1\text{-}114}$, masking thus $^{FIRE}$mate. The structure of apo PYP was generated from the crystal structure (PDB: 1NWZ). The model of fluorogen-bound pFAST was generated by homology modeling and molecular dynamics in ref.[1]. The model of caged$^{FIRE}$tag was predicted using Alphafold[2].

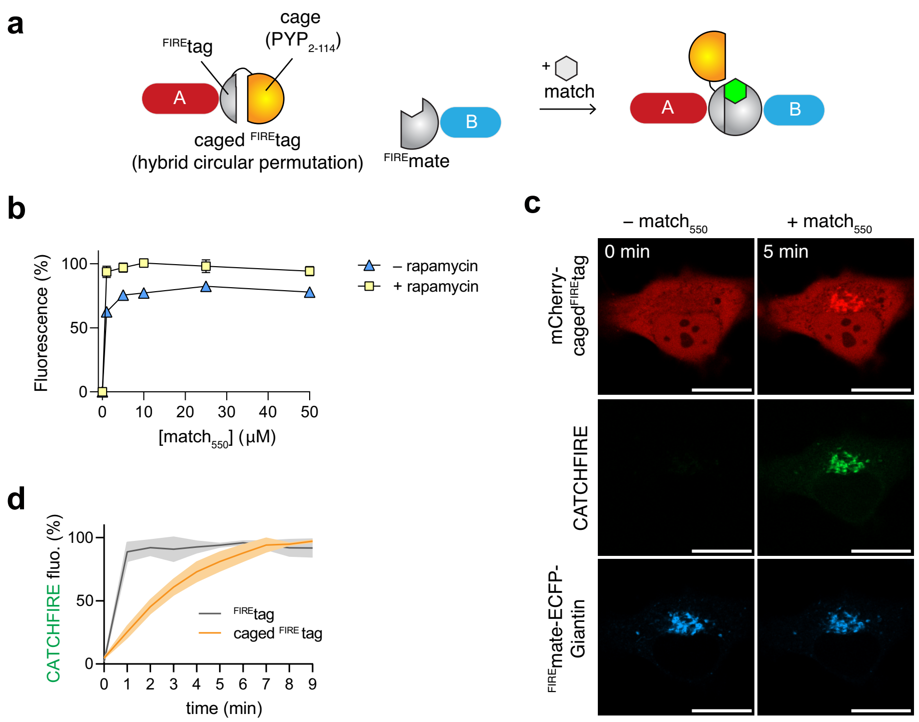


**Figure S8. Caging $^{FIRE}$tag avoids undesirable residual self-association. a** Design of caged$^{FIRE}$tag. **b** Normalized average fluorescence of about 50,000 HEK293T cells co-expressing the FK506-binding protein (FKBP) fused to caged$^{FIRE}$tag and the FKBP-rapamycin-binding domain of mammalian target of rapamycin (FRB) fused to $^{FIRE}$mate treated without or with 500 nM of rapamycin, and with 1, 5, 10, 25 or 50 µM of match$_{550}$. Individual cell fluorescence was analyzed by flow cytometry. Data represent the mean values ± SD of three independent experiments. **c,d** HeLa cells co-expressing mCherry-caged$^{FIRE}$tag and $^{FIRE}$mate-ECFP-Giantin were treated with 10 µM match$_{550}$, and imaged by time-lapse confocal microscopy. **c** Representative confocal micrographs of cells before (0 min) and after (9 min) addition of match$_{550}$ (see also **Movie S16**). Experiments were repeated three times with similar results. Red: ex/em 561/606-675 nm; green: ex/em 488/508-570 nm; cyan: ex/em 445/455-499 nm. Scale bars are 20 µm. **c** Temporal evolution of the CATCHFIRE signal. Data represents the mean values ± SD of 7 cells from three independent experiments.

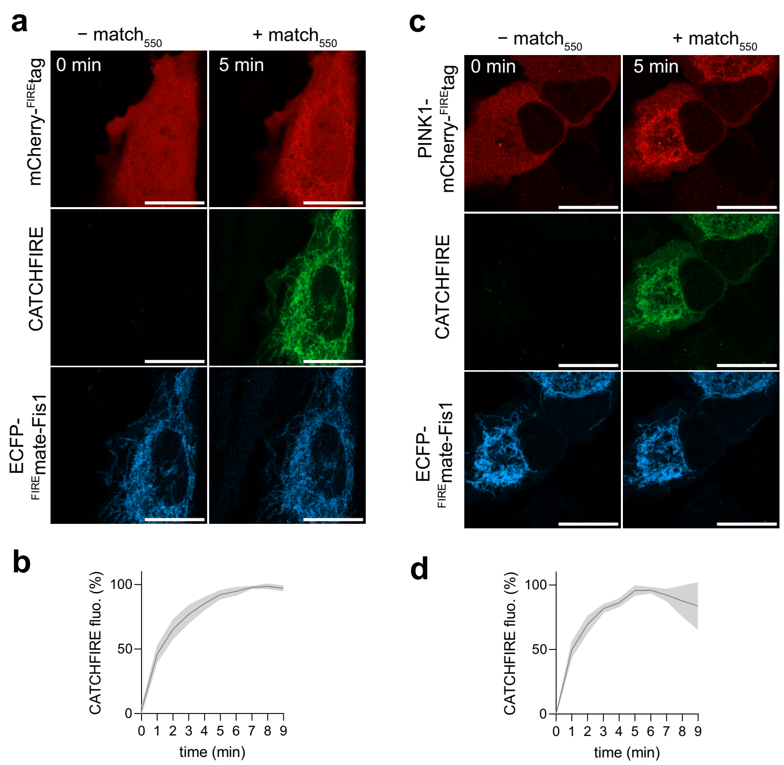


**Figure S9. Fluorogenic induced recruitment of PINK1 to mitochondria.** HeLa cells co-expressing ECFP-$^{FIRE}$mate-Fis1 and either **a,b** mCherry-$^{FIRE}$tag or **c,d** PINK1-mCherry-$^{FIRE}$tag were treated without or with $match_{550}$, and imaged by time-lapse confocal microscopy. **a,c** Representative confocal micrographs of cells before (0 min) and after (5 min) addition of the fluorogenic inducer of proximity. Experiments were repeated three times with similar results. **b,d** Temporal evolution of the recruitment. Data represents the mean values ± SD of 8 cells (**a,b**) and 16 cells (**c,d**) from three independent experiments. Red: ex/em 561/606-675 nm; green: ex/em 488/508-570 nm; cyan: ex/em 445/455-499 nm).

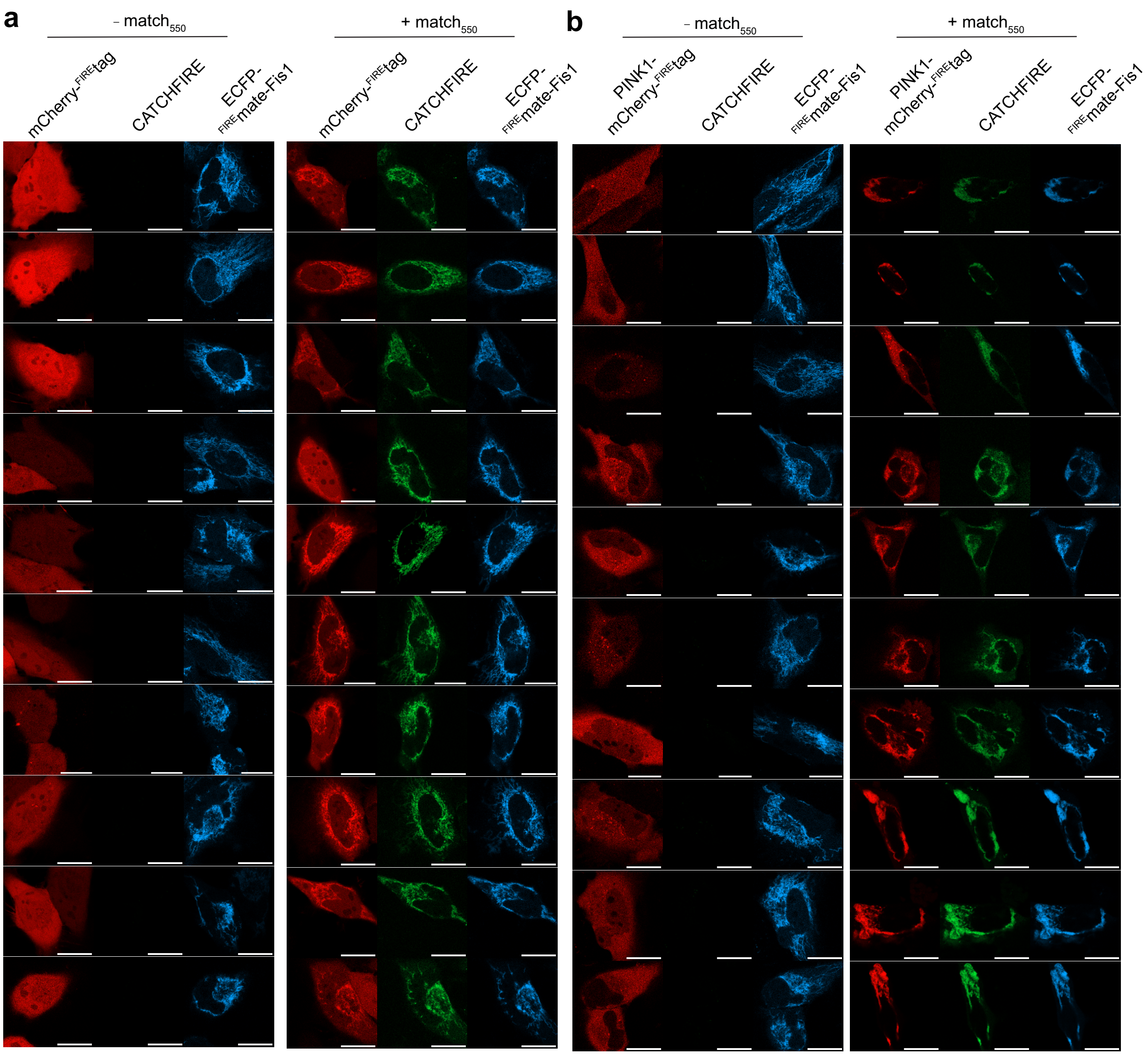


**Figure S10. Fluorogenic induced recruitment of PINK1 to mitochondria induces mitophagy.** HeLa cells co-expressing ECFP-$^{FIRE}$mate-Fis1 and either **a** mCherry-$^{FIRE}$tag (negative control) or **b** PINK1-mCherry-$^{FIRE}$tag were treated without or with match$_{550}$ for 2 h. Experiments were repeated three times with similar results. Representative micrographs are shown. Red: ex/em 561/606-675 nm; green: ex/em 488/508-570 nm; cyan: ex/em 445/455-499 nm).

### Table S1. DNA sequences

| Vector | ORF | ORF sequence |
|---|---|---|
| | $^{\text{FIRE}}$tag | ggtgacagatattgggtctttgtgaaacgggtg |
| | $^{\text{FIRE}}$mate | atggagcatgttgcctttggcagtgaggacatcgagaacactctggccaatatggacgacgaacaactggataggttggcctttggcgta<br>attcagctcgatggtgacgggaatatcctgctgtacaatgctgctgaaggggacatcactggcagagatcccaaacaggtgattgggaag<br>aacttcttcaaggatgttgcacctggaacggatactcccgagttttacggcaaattcaaggaaggcgcagcgtcagggaatctgaacacc<br>atgttcgaatggacgataccgacaagcaggggaccaaccaaggtcaaggtgcacttgaagaaagccctttcc |
| | caged $^{\text{FIRE}}$tag | ggtgacagatattgggtctttgtgaaacgggtggaaaacctgtattttcagggcggctccggcggcgaacatgttgcctttggttcggaa<br>gatatcgagaacactctggcgaaaatggacgacggtcaactggatggcttggcttttggtgccattcaactggatggcgatggcaatatc<br>ctccagtataatgcggctgaaggggacattaccggacgtgatccgaaacaggtcattgggaagaacttcttcaaagacgtagcaccaggt<br>acggatagtccggaattctacggcaagttcaaagaaggagttgcatctggcaatctgaacacgatgtttgagtatacctttgattaccag<br>atgacacctaccaaagtgaaagtccacatgaagaaagcgttatcc |
| pAG1159 | TOM20(1-34)-ECFP-$^{\text{FIRE}}$mate | atggtgggtcggaacagcgccatcgccgcgggcgtgtgcggtgccctcttcatagggtactgcatctactttgaccgcaaaagacgaagt<br>gaccccaacttcggatccagtgctggtggtagtgctggtggtatggtgagcaagggcgaggagctgttcaccggggtggtgcccatcctg<br>gtcgagctggacggcgacgtaaacggccacaagttcagcgtgtccggcgagggcgagggcgatgccacctacggcaagctgaccctgaag<br>ttcatctgcaccaccggcaagctgcccgtgccctggcccaccctcgtgaccaccctgacctggggcgtgcagtgcttcagccgctacccc<br>gaccacatgaagcagcacgacttcttcaagtccgccatgcccgaaggctacgtccaggagcgcaccatcttcttcaaggacgacggcaac<br>tacaagacccgcgccgaggtgaagttcgagggcgacaccctggtgaaccgcatcgagctgaagggcatcgacttcaaggaggacggcaac<br>atcctggggcacaagctggagtacaactacatcagccacaacgtctatatcaccgccgacaagcagaagaacggcatcaaggccaacttc<br>aagatccgccacaacatcgaggacggcagcgtgcagctcgccgaccactaccagcagaacacccccatcggcgacggccccgtgctgctg<br>cccgacaaccactacctgagcacccagtccgccctgagcaaagaccccaacgagaagcgcgatcacatggtcctgctggagttcgtgacc<br>gccgccgggatcactctcggcatggacgagctgtacaagggagcaagtggaatggagcatgttgcctttggcagtgaggacatcgagaac<br>actctggccaatatggacgacgaacaactggataggttggcctttggcgtaattcagctcgatggtgacgggaatatcctgctgtacaat<br>gctgctgaaggggacatcactggcagagatcccaaacaggtgattgggaagaacttcttcaaggatgttgcacctggaacggatactccc<br>gagttttacggcaaattcaaggaaggcgcagcgtcagggaatctgaacaccatgttcgaatggacgataccgacaagcaggggaccaacc<br>aaggtcaaggtgcacttgaagaaagccctttcc |
| pAG1160 | mCherry-$^{\text{FIRE}}$tag | atggtgagcaagggcgaggaggataacatggccatcatcaaggagttcatgcgcttcaaggtgcacatggagggctccgtgaacggccac<br>gagttcgagatcgagggcgagggcgagggccgcccctacgagggcacccagaccgccaagctgaaggtgaccaagggtggccccctgccc<br>ttcgcctgggacatcctgtcccctcagttcatgtacggctccaaggcctacgtgaagcaccccgccgacatccccgactacttgaagctg<br>tccttccccgagggcttcaagtgggagcgcgtgatgaacttcgaggacggcggcgtggtgaccgtgacccaggactcctccctgcaggac<br>ggcgagttcatctacaaggtgaagctgcgcggcaccaacttcccctccgacggccccgtaatgcagaagaagaccatgggctgggaggcc<br>tcctccgagcggatgtaccccgaggacggcgccctgaagggcgagatcaagcagaggctgaagctgaaggacggcggccactacgacgct<br>gaggtcaagaccacctacaaggccaagaagcccgtgcagctgcccggcgcctacaacgtcaacatcaagttggacatcacctcccacaac<br>gaggactacaccatcgtggaacagtacgaacgcgccgagggccgccactccaccggcggcatggacgagctgtacaagtctagaggcggc<br>ggctccggtgacagatattgggtctttgtgaaacgggtg |
| pAG1344 | EGFP- $^{\text{FIRE}}$tag | atggtgagcaagggcgaggagctgttcaccggggtggtgcccatcctggtcgagctggacggcgacgtaaacggccacaagttcagcgtg<br>tccggcgagggcgagggcgatgccacctacggcaagctgaccctgaagttcatctgcaccaccggcaagctgcccgtgccctggcccacc<br>ctcgtgaccaccctgacctacggcgtgcagtgcttcagccgctaccccgaccacatgaagcagcacgacttcttcaagtccgccatgccc<br>gaaggctacgtccaggagcgcaccatcttcttcaaggacgacggcaactacaagacccgcgccgaggtgaagttcgagggcgacaccctg<br>gtgaaccgcatcgagctgaagggcatcgacttcaaggaggacggcaacatcctggggcacaagctggagtacaactacaacagccacaac<br>gtctatatcatggccgacaagcagaagaacggcatcaaggtgaacttcaagatccgccacaacatcgaggacggcagcgtgcagctcgcc<br>gaccactaccagcagaacacccccatcggcgacggccccgtgctgctgcccgacaaccactacctgagcacccagtccgccctgagcaaa<br>gaccccaacgagaagcgcgatcacatggtcctgctggagttcgtgaccgccgccgggatcactctcggcatggacgagctgtacaagtct<br>agaggcggcggctccggtgacagatattgggtctttgtgaaacgggtg |
| pAG1171 | $^{\text{FIRE}}$mate-ECFP-Giantin(3131-3259) | atggagcatgttgcctttggcagtgaggacatcgagaacactctggccaatatggacgacgaacaactggataggttggcctttggcgta<br>attcagctcgatggtgacgggaatatcctgctgtacaatgctgctgaaggggacatcactggcagagatcccaaacaggtgattgggaag<br>aacttcttcaaggatgttgcacctggaacggatactcccgagttttacggcaaattcaaggaaggcgcagcgtcagggaatctgaacacc<br>atgttcgaatggacgataccgacaagcaggggaccaaccaaggtcaaggtgcacttgaagaaagccctttccggtggtgctagtggtggt<br>gctagtatggtgagcaagggcgaggagctgttcaccggggtggtgcccatcctggtcgagctggacggcgacgtaaacggccacaagttc<br>agcgtgtccggcgagggcgagggcgatgccacctacggcaagctgaccctgaagttcatctgcaccaccggcaagctgcccgtgccctgg<br>cccaccctcgtgaccaccctgacctggggcgtgcagtgcttcagccgctaccccgaccacatgaagcagcacgacttcttcaagtccgcc<br>atgcccgaaggctacgtccaggagcgcaccatcttcttcaaggacgacggcaactacaagacccgcgccgaggtgaagttcgagggcgac<br>accctggtgaaccgcatcgagctgaagggcatcgacttcaaggaggacggcaacatcctggggcacaagctggagtacaactacatcagc<br>cacaacgtctatatcaccgccgacaagcagaagaacggcatcaaggccaacttcaagatccgccacaacatcgaggacggcagcgtgcag<br>ctcgccgaccactaccagcagaacacccccatcggcgacggccccgtgctgctgcccgacaaccactacctgagcacccagtccgccctg<br>agcaaagaccccaacgagaagcgcgatcacatggtcctgctggagttcgtgaccgccgccgggatcactctcggcatggacgagctgtac<br>aagtccggactcagatctcgaggagaaccgcagcaaagcttttctgaagctcagcagcagctatgcaacaccagacaggaagtgaatgaa<br>ttaaggaagctgctggaagaagaacgagaccaaagagtggctgctgagaatgctctctctgtggccgaggagcagatcagacggttagag<br>cacagtgaatgggactcttcccggactcctatcattggctcctgtggcactcaggagcaggcactgttaatagatcttacaagcaacagt<br>tgtcgaaggacccggagtggcgttggatggaagcgagtcctgcgttcactctgtcattcacggacccgagtgccacttctagcagccatc<br>tactttctaatgattcatgtcctgctcattctgtgttttacgggccatcta |
| pAG1169 | $^{\text{FIRE}}$mate-ECFP-Cb5(100-134) | atggagcatgttgcctttggcagtgaggacatcgagaacactctggccaatatggacgacgaacaactggataggttggcctttggcgta<br>attcagctcgatggtgacgggaatatcctgctgtacaatgctgctgaaggggacatcactggcagagatcccaaacaggtgattgggaag<br>aacttcttcaaggatgttgcacctggaacggatactcccgagttttacggcaaattcaaggaaggcgcagcgtcagggaatctgaacacc<br>atgttcgaatggacgataccgacaagcaggggaccaaccaaggtcaaggtgcacttgaagaaagccctttccggtggtgctagtggtggt<br>gctagtatggtgagcaagggcgaggagctgttcaccggggtggtgcccatcctggtcgagctggacggcgacgtaaacggccacaagttc<br>agcgtgtccggcgagggcgagggcgatgccacctacggcaagctgaccctgaagttcatctgcaccaccggcaagctgcccgtgccctgg<br>cccaccctcgtgaccaccctgacctggggcgtgcagtgcttcagccgctaccccgaccacatgaagcagcacgacttcttcaagtccgcc<br>atgcccgaaggctacgtccaggagcgcaccatcttcttcaaggacgacggcaactacaagacccgcgccgaggtgaagttcgagggcgac<br>accctggtgaaccgcatcgagctgaagggcatcgacttcaaggaggacggcaacatcctggggcacaagctggagtacaactacatcagc<br>cacaacgtctatatcaccgccgacaagcagaagaacggcatcaaggccaacttcaagatccgccacaacatcgaggacggcagcgtgcag<br>ctcgccgaccactaccagcagaacacccccatcggcgacggccccgtgctgctgcccgacaaccactacctgagcacccagtccgccctg<br>agcaaagaccccaacgagaagcgcgatcacatggtcctgctggagttcgtgaccgccgccgggatcactctcggcatggacgagctgtac<br>aagtccggactcagatctatcaccaccgtggagtccaactcctcctggtggaccaactgggtgatccccgccatctccgccctggtggtg<br>gccctgatgtaccgcctatacatggccgaggac |

| | | |
|---|---|---|
| pAG1186 | Lyn11-ECFP-<br>$^{FIRE}$mate | atgggctgcatcaagtccaagggcaaggactccgccggtagtggtggtatggtgagcaagggcgaggagctgttcaccggggtggtgcccat<br>cctggtcgagctggacggcgacgtaaacggccacaagttcagcgtgtccggcgagggcgagggcgatgccacctacggcaagctgaccctga<br>agttcatctgcaccaccggcaagctgcccgtgccctggcccaccctcgtgaccaccctgacctggggcgtgcagtgcttcagccgctacccc<br>gaccacatgaagcagcacgacttcttcaagtccgccatgcccgaaggctacgtccaggagcgcaccatcttcttcaaggacgacggcaacta<br>caagacccgcgccgaggtgaagttcgagggcgacaccctggtgaaccgcatcgagctgaagggcatcgacttcaaggaggacggcaacatcc<br>tggggcacaagctggagtacaactacatcagccacaacgtctatatcaccgccgacaagcagaagaacggcatcaaggccaacttcaagatc<br>cgccacaacatcgaggacggcagcgtgcagctcgccgaccactaccagcagaacacccccatcggcgacggccccgtgctgctgcccgacaa<br>ccactacctgagcacccagtccgccctgagcaaagaccccaacgagaagcgcgatcacatggtcctgctggagttcgtgaccgccgccggga<br>tcactctcggcatggacgagctgtacaagggagcaagtggaatggagcatgttgcctttggcagtgaggacatcgagaacactctggccaat<br>atggacgacgaacaactggataggttggcctttggcgtaattcagctcgatggtgacgggaatatcctgctgtacaatgctgctgaaggga<br>catcactggcagagatcccaaacaggtgattgggaagaacttcttcaaggatgttgcacctggaacggatactcccgagttttacggcaaat<br>tcaaggaaggcgcagcgtcagggaatctgaacaccatgttcgaatggacgataccgacaagcaggggaccaaccaaggtcaaggtgcacttg<br>aagaaagccctttcc |
| pAG1243 | NES-ECFP-<br>$^{FIRE}$mate | atgttagccttgaaattagcaggtcttgatatcgggagcggtagtgctggtggtatggtgagcaagggcgaggagctgttcaccggggtggt<br>gcccatcctggtcgagctggacggcgacgtaaacggccacaagttcagcgtgtccggcgagggcgagggcgatgccacctacggcaagctga<br>ccctgaagttcatctgcaccaccggcaagctgcccgtgccctggcccaccctcgtgaccaccctgacctggggcgtgcagtgcttcagccgc<br>taccccgaccacatgaagcagcacgacttcttcaagtccgccatgcccgaaggctacgtccaggagcgcaccatcttcttcaaggacgacgg<br>caactacaagacccgcgccgaggtgaagttcgagggcgacaccctggtgaaccgcatcgagctgaagggcatcgacttcaaggaggacggca<br>acatcctggggcacaagctggagtacaactacatcagccacaacgtctatatcaccgccgacaagcagaagaacggcatcaaggccaacttc<br>aagatccgccacaacatcgaggacggcagcgtgcagctcgccgaccactaccagcagaacacccccatcggcgacggccccgtgctgctgcc<br>cgacaaccactacctgagcacccagtccgccctgagcaaagaccccaacgagaagcgcgatcacatggtcctgctggagttcgtgaccgccg<br>ccgggatcactctcggcatggacgagctgtacaagggagcaagtggaatggagcatgttgcctttggcagtgaggacatcgagaacactctg<br>gccaatatggacgacgaacaactggataggttggcctttggcgtaattcagctcgatggtgacgggaatatcctgctgtacaatgctgctga<br>aggggacatcactggcagagatcccaaacaggtgattgggaagaacttcttcaaggatgttgcacctggaacggatactcccgagttttacg<br>gcaaattcaaggaaggcgcagcgtcagggaatctgaacaccatgttcgaatggacgataccgacaagcaggggaccaaccaaggtcaaggtg<br>cacttgaagaaagccctttcc |
| pAG1242 | NLS-mCherry-<br>$^{FIRE}$tag | atgccaaaaaagaaaagaaaagtttccggaggaggcggcatggtgagcaagggcgaggaggataacatggccatcatcaaggagttcatgcg<br>cttcaaggtgcacatggagggctccgtgaacggccacgagttcgagatcgagggcgagggcgagggccgcccctacgagggcacccagaccg<br>ccaagctgaaggtgaccaagggtggccccctgcccttcgcctgggacatcctgtcccctcagttcatgtacggctccaaggcctacgtgaag<br>caccccgccgacatccccgactacttgaagctgtccttccccgagggcttcaagtgggagcgcgtgatgaacttcgaggacggcggcgtggt<br>gaccgtgacccaggactcctccctgcaggacggcgagttcatctacaaggtgaagctgcgcggcaccaacttcccctccgacggccccgtaa<br>tgcagaagaagaccatgggctgggaggcctcctccgagcggatgtaccccgaggacggcgccctgaagggcgagatcaagcagaggctgaag<br>ctgaaggacggcggccactacgacgctgaggtcaagaccacctacaaggccaagaagcccgtgcagctgcccggcgcctacaacgtcaacat<br>caagttggacatcacctcccacaacgaggactacaccatcgtggaacagtacgaacgcgccgagggccgccactccaccggcggcatggacg<br>agctgtacaagtctagaggcggcggctccggtgacagatattgggtctttgtgaaacgggtg |
| pAG1247 | H2B-ECFP-<br>$^{FIRE}$mate | atgcccgaacctgcgaagtcagcgcccgctcccaaaaaaggctctaaaaagctgtcgccaagacccagaagaaggggataagaaaaggcg<br>taagaccaggaaagagagttacgccatttacgtgtacaaagtactaaaacaagtccacccggacactggcatctcctcaaaggcgatgggca<br>ttatgaactcatttgtaaacgacatcttcgagcgcatcgccggagaagcgtcgcgcctggcgcattacaacaagcgctccactatcacatcc<br>cgggagatccagacggccgtgcgcctgctcctgcccggagaactggccaaacacgctgtgtctgagggcacaaaggccgtgaccaagtacac<br>cagctccaagggtagtgctggtggtatggtgagcaagggcgaggagctgttcaccggggtggtgcccatcctggtcgagctggacggcgacg<br>taaacggccacaagttcagcgtgtccggcgagggcgagggcgatgccacctacggcaagctgaccctgaagttcatctgcaccaccggcaag<br>ctgcccgtgccctggcccaccctcgtgaccaccctgacctggggcgtgcagtgcttcagccgctaccccgaccacatgaagcagcacgactt<br>cttcaagtccgccatgcccgaaggctacgtccaggagcgcaccatcttcttcaaggacgacggcaactacaagacccgcgccgaggtgaagt<br>tcgagggcgacaccctggtgaaccgcatcgagctgaagggcatcgacttcaaggaggacggcaacatcctggggcacaagctggagtacaac<br>tacatcagccacaacgtctatatcaccgccgacaagcagaagaacggcatcaaggccaacttcaagatccgccacaacatcgaggacggcag<br>cgtgcagctcgccgaccactaccagcagaacacccccatcggcgacggccccgtgctgctgcccgacaaccactacctgagcacccagtccg<br>ccctgagcaaagaccccaacgagaagcgcgatcacatggtcctgctggagttcgtgaccgccgccgggatcactctcggcatggacgagctg<br>tacaagggagcaagtggaatggagcatgttgcctttggcagtgaggacatcgagaacactctggccaatatggacgacgaacaactggatag<br>gttggcctttggcgtaattcagctcgatggtgacgggaatatcctgctgtacaatgctgctgaaggggacatcactggcagagatcccaaac<br>aggtgattgggaagaacttcttcaaggatgttgcacctggaacggatactcccgagttttacggcaaattcaaggaaggcgcagcgtcaggg<br>aatctgaacaccatgttcgaatggacgataccgacaagcaggggaccaaccaaggtcaaggtgcacttgaagaaagccctttcc |
| pAG1280 | NLS-mCherry-<br>$^{FIRE}$tag- NES | atgccaaaaaagaaaagaaaagtttccggaggaggcggcatggtgagcaagggcgaggaggataacatggccatcatcaaggagttcatgcg<br>cttcaaggtgcacatggagggctccgtgaacggccacgagttcgagatcgagggcgagggcgagggccgcccctacgagggcacccagaccg<br>ccaagctgaaggtgaccaagggtggccccctgcccttcgcctgggacatcctgtcccctcagttcatgtacggctccaaggcctacgtgaag<br>caccccgccgacatccccgactacttgaagctgtccttccccgagggcttcaagtgggagcgcgtgatgaacttcgaggacggcggcgtggt<br>gaccgtgacccaggactcctccctgcaggacggcgagttcatctacaaggtgaagctgcgcggcaccaacttcccctccgacggccccgtaa<br>tgcagaagaagaccatgggctgggaggcctcctccgagcggatgtaccccgaggacggcgccctgaagggcgagatcaagcagaggctgaag<br>ctgaaggacggcggccactacgacgctgaggtcaagaccacctacaaggccaagaagcccgtgcagctgcccggcgcctacaacgtcaacat<br>caagttggacatcacctcccacaacgaggactacaccatcgtggaacagtacgaacgcgccgagggccgccactccaccggcggcatggacg<br>agctgtacaagtctagaggcggcggctccggtgacagatattgggtctttgtgaaacgggtgggctccggcggcgatgaggtggatggctcc<br>ggcggcttagccttgaaattagcaggtcttgatatcgggagc |
| pAG1237 | $^{FIRE}$mate-ECFP-<br>Fis1 | atggagcatgttgcctttggcagtgaggacatcgagaacactctggccaatatggacgacgaacaactggataggttggcctttggcgtaat<br>tcagctcgatggtgacgggaatatcctgctgtacaatgctgctgaaggggacatcactggcagagatcccaaacaggtgattgggaagaact<br>tcttcaaggatgttgcacctggaacggatactcccgagttttacggcaaattcaaggaaggcgcagcgtcagggaatctgaacaccatgttc<br>gaatggacgataccgacaagcaggggaccaaccaaggtcaaggtgcacttgaagaaagccctttccggtggtgctagtggtggtgctagtat<br>ggtgagcaagggcgaggagctgttcaccggggtggtgcccatcctggtcgagctggacggcgacgtaaacggccacaagttcagcgtgtccg<br>gcgagggcgagggcgatgccacctacggcaagctgaccctgaagttcatctgcaccaccggcaagctgcccgtgccctggcccaccctcgtg<br>accaccctgacctggggcgtgcagtgcttcagccgctaccccgaccacatgaagcagcacgacttcttcaagtccgccatgcccgaaggcta<br>cgtccaggagcgcaccatcttcttcaaggacgacggcaactacaagacccgcgccgaggtgaagttcgagggcgacaccctggtgaaccgca<br>tcgagctgaagggcatcgacttcaaggaggacggcaacatcctggggcacaagctggagtacaactacatcagccacaacgtctatatcacc<br>gccgacaagcagaagaacggcatcaaggccaacttcaagatccgccacaacatcgaggacggcagcgtgcagctcgccgaccactaccagca<br>gaacacccccatcggcgacggccccgtgctgctgcccgacaaccactacctgagcacccagtccgccctgagcaaagaccccaacgagaagc<br>gcgatcacatggtcctgctggagttcgtgaccgccgccgggatcactctcggcatggacgagctgtacaagtgtacaagtctagaggcggcg<br>gcttacgtccgcgggttgctgcagacagagccccagaacaaccaggccaaggaactggagcggctcattgacaaggccatgaagaaagatgg<br>actcgtgggcatggccatcgtgggaggcatggccctgggtgtggcgggactggccggactcatcggacttgctgtgtccaagtccaaatcc |

| | | |
|---|---|---|
| pAG1238 | PINK1-mCherry-$^{\text{FIRE}}$tag | atgatcgaggaaaaacaggcggagagccggcgggcggtctcggcctgtcaggagatccaggcaatttttacccagaaaagcaagccggggc<br>ctgacccgttggacacgagacgcttgcagggctttcggctggaggagtatctgatagggcagtccattggtaagggctgcagtgctgctgt<br>gtatgaagccaccatgcctacattgccccagaacctggaggtgacaaagagcaccgggttgcttccagggagaggcccaggtaccagtgca<br>ccaggagaagggcaggagcgagctccgggggcccctgccttccccttggccatcaagatgatgtggaacatctcggcaggttcctccagcg<br>aagccatcttgaacacaatgagccaggagctggtcccagcgagccgagtggccttggctggggagtatggagcagtcacttacagaaaatc<br>caagagaggtcccaagcaactagcccctcaccccaacatcatccgggttctccgcgccttcacctcttccgtgccgctgctgccaggggcc<br>ctggtcgactaccctgatgtgctgccctcacgcctccaccctgaaggcctgggccatggccggacgctgttcctcgttatgaagaactatc<br>cctgtaccctgcgccagtacctttgtgtgaacacacccagcccccgcctcgccgccatgatgctgctgcagctgctggaaggcgtggacca<br>tctggttcaacagggcatcgcgcacagagacctgaaatccgacaacatccttgtggagctggacccagacggctgccccctggctggtgatc<br>gcagattttggctgctgcctggctgatgagagcatcggcctgcagttgcccttcagcagctggtacgtggatcggggcggaaacggctgtc<br>tgatggccccagaggtgtccacggcccgtcctggccccagggcagtgattgactacagcaaggctgatgcctgggcagtgggagccatcgc<br>ctatgaaatcttcgggcttgtcaatcccttctacggccagggcaaggcccaccttgaaagccgcagctaccaagaggctcagctacctgca<br>ctgcccgagtcagtgcctccagacgtgagacagttggtgagggcactgctccagcgagaggccagcaagagaccatctgcccgagtagccg<br>caaatgtgcttcatctaagcctctggggtgaacatattctagccctgaagaatctgaagttagacaagatggttggctggctcctccaaca<br>atcggccgccactttgttggccaacaggctcacagagaagtgttgtgtggaaacaaaaatgaagatgctctttctggctaacctggagtgt<br>gaaacgctctgccaggcagccctcctcctctgctcatggagggcagccctgagcggcgggggaggctccggagggatggtgagcaagggcg<br>aggaggataacatggccatcatcaaggagttcatgcgcttcaaggtgcacatggagggctccgtgaacggccacgagttcgagatcgaggg<br>cgagggcgagggccgcccctacgagggcacccagaccgccaagctgaaggtgaccaagggtggccccctgcccttcgcctgggacatcctg<br>tcccctcagttcatgtacggctccaaggcctacgtgaagcaccccgccgacatccccgactacttgaagctgtccttccccgagggcttca<br>agtgggagcgcgtgatgaacttcgaggacggcggcgtggtgaccgtgacccaggactcctccctgcaggacggcgagttcatctacaaggt<br>gaagctgcgcggcaccaacttcccctccgacggccccgtaatgcagaagaagaccatgggctgggaggcctcctccgagcggatgtacccc<br>gaggacggcgccctgaagggcgagatcaagcagaggctgaagctgaaggacggcggccactacgacgctgaggtcaagaccacctacaagg<br>ccaagaagcccgtgcagctgcccggcgcctacaacgtcaacatcaagttggacatcacctcccacaacgaggactacaccatcgtggaaca<br>gtacgaacgcgccgagggccgccactccaccggcggcatggacgagctgtacaagtctagaggcggcggctccggtgacagatattgggtc<br>tttgtgaaacgggtg |
| pAG1244 | p65-mCherry-$^{\text{FIRE}}$tag | atggacgaactgttcccccctcatcttcccggcagagccagcccaggcctctggcccctatgtggagatcattgagcagcccaagcagcggg<br>gcatgcgcttccgctacaagtgcgagggcgctccgcgggcagcatcccaggcgagaggagcacagataccaccaagacccaccccaccat<br>caagatcaatggctacacaggaccagggacagtgcgcatctccctggtcaccaaggaccctcctcaccggcctcacccccacgagcttgta<br>ggaaaggactgccgggatggcttctatgaggctgagctctgcccggaccgctgcatccacagtttccagaacctgggaatccagtgtgtga<br>agaagcgggacctggagcaggctatcagtcagcgcatccagaccaacaacaaccccttccaagttcctatagaagagcagcgtggggacta<br>cgacctgaatgctgtgcggctctgcttccaggtgacagtgcgggacccatcaggcaggcccctccgcctgccgcctgtcctttctcatccc<br>atctttgacaatcgtgcccccaacactgccgagctcaagatctgccgagtgaaccgaaactctggcagctgcctcggtggggatgagatct<br>tcctactgtgtgacaaggtgcagaaagaggacattgaggtgtatttcacgggaccaggctgggaggcccgaggctccttttcgcaagctga<br>tgtgcaccgacaagtggccattgtgttccggacccctccctacgcagaccccagcctgcaggctcctgtgcgtgtctccatgcagctgcgg<br>cggccttccgaccgggagctcagtgagcccatggaattccagtacctgccagatacagacgatcgtcaccggattgaggagaaacgtaaaa<br>ggacatatgagaccttcaagagcatcatgaagaagagtcctttcagcggacccaccgacccccggcctccacctcgacgcattgctgtgcc<br>ttcccgcagctcagcttctgtccccaagccagcacccagccctatccctttacgtcatccctgagcaccatcaactatgatgagtttccc<br>accatggtgtttccttctgggcagatcagccaggcctcggccttggccccggcccctccccaagtcctgccccaggctccagcccctgccc<br>ctgctccagccatggtatcagctctggcccaggccccagcccctgtcccagtcctagccccaggccctcctcaggctgtggccccacctgc<br>ccccaagcccacccaggctggggaaggaacgctgtcagaggccctgctgcagctgcagtttgatgatgaagacctgggggccttgcttggc<br>aacagcacagacccagctgtgttcacagacctggcatccgtcgacaactccgagtttcagcagctgctgaaccagggcatacctgtggccc<br>cccacacaactgagcccatgctgatggagtaccctgaggctataactcgcctagtgacaggggcccagaggcccccgacccagctcctgc<br>tccactggggcccgggctccccaatggcctcctttcaggagatgaagacttctcctccattgcggacatggacttctcagccctgctg<br>agtcagatcagctcctccggaggaggcggcatggtgagcaagggcgaggaggataacatggccatcatcaaggagttcatgcgcttcaagg<br>tgcacatggagggctccgtgaacggccacgagttcgagatcgagggcgagggcgagggccgcccctacgagggcacccagaccgccaagct<br>gaaggtgaccaagggtggccccctgcccttcgcctgggacatcctgtcccctcagttcatgtacggctccaaggcctacgtgaagcacccc<br>gccgacatccccgactacttgaagctgtccttccccgagggcttcaagtgggagcgcgtgatgaacttcgaggacggcggcgtggtgaccg<br>tgacccaggactcctccctgcaggacggcgagttcatctacaaggtgaagctgcgcggcaccaacttcccctccgacggccccgtaatgca<br>gaagaagaccatgggctgggaggcctcctccgagcggatgtaccccgaggacggcgccctgaagggcgagatcaagcagaggctgaagctg<br>aaggacggcggccactacgacgctgaggtcaagaccacctacaaggccaagaagcccgtgcagctgcccggcgcctacaacgtcaacatca<br>agttggacatcacctcccacaacgaggactacaccatcgtggaacagtacgaacgcgccgagggccgccactccaccggcggcatggacga<br>gctgtacaagtctagaggcggcggctccggtgacagatattgggtctttgtgaaacgggtg |
| pAG1345 | mCherry- $^{\text{FIRE}}$tag-**DEVDG**-MoA(2880-2996) | atggtgagcaagggcgaggaggataacatggccatcatcaaggagttcatgcgcttcaaggtgcacatggagggctccgtgaacggccacg<br>agttcgagatcgagggcgagggcgagggccgcccctacgagggcacccagaccgccaagctgaaggtgaccaagggtggccccctgccctt<br>cgcctgggacatcctgtcccctcagttcatgtacggctccaaggcctacgtgaagcaccccgccgacatccccgactacttgaagctgtcc<br>ttccccgagggcttcaagtgggagcgcgtgatgaacttcgaggacggcggcgtggtgaccgtgacccaggactcctccctgcaggacggcg<br>agttcatctacaaggtgaagctgcgcggcaccaacttcccctccgacggccccgtaatgcagaagaagaccatgggctgggaggcctcctc<br>cgagcggatgtaccccgaggacggcgccctgaagggcgagatcaagcagaggctgaagctgaaggacggcggccactacgacgctgaggtc<br>aagaccacctacaaggccaagaagcccgtgcagctgcccggcgcctacaacgtcaacatcaagttggacatcacctcccacaacgaggact<br>acaccatcgtggaacagtacgaacgcgccgagggccgccactccaccggcggcatggacgagctgtacaagtctagaggcggcggctccgg<br>tgacagatattgggtctttgtgaaacgggtgggctccggcggc**GATGAGGTGGATGGC**tccggcggcagtgctggtggtagtgctggtggt<br>agtgctggtggtactggtggtcctcgagctcaagcttcgaattcgagtgctggtggtttctgggaaaggaacctgccctctgtttctggcc<br>tgctgaagatcattggattttccacatcagtaactgccctggggtttgtgctgtacaaatacaagctcctgccacggtcttga |
| pAG1188 | mCherry-caged$^{\text{FIRE}}$tag | atggtgagcaagggcgaggaggataacatggccatcatcaaggagttcatgcgcttcaaggtgcacatggagggctccgtgaacggccacg<br>agttcgagatcgagggcgagggcgagggccgcccctacgagggcacccagaccgccaagctgaaggtgaccaagggtggccccctgccctt<br>cgcctgggacatcctgtcccctcagttcatgtacggctccaaggcctacgtgaagcaccccgccgacatccccgactacttgaagctgtcc<br>ttccccgagggcttcaagtgggagcgcgtgatgaacttcgaggacggcggcgtggtgaccgtgacccaggactcctccctgcaggacggcg<br>agttcatctacaaggtgaagctgcgcggcaccaacttcccctccgacggccccgtaatgcagaagaagaccatgggctgggaggcctcctc<br>cgagcggatgtaccccgaggacggcgccctgaagggcgagatcaagcagaggctgaagctgaaggacggcggccactacgacgctgaggtc<br>aagaccacctacaaggccaagaagcccgtgcagctgcccggcgcctacaacgtcaacatcaagttggacatcacctcccacaacgaggact<br>acaccatcgtggaacagtacgaacgcgccgagggccgccactccaccggcggcatggacgagctgtacaagtctagaggcggcggctccgg<br>tgacagatattgggtctttgtgaaacgggtggaaaacctgtattttcagggcggctccggcggcgaacatgttgcctttggttcggaagat<br>atcgagaacactctggcgaaaatggacgacggtcaactggatggcttggcttttggtgccattcaactggatggcgatggcaatatcctcc<br>agtataatgcggctgaaggggacattaccggacgtgatccgaaacaggtcattgggaagaacttcttcaaagacgtagcaccaggtacgga<br>tagtccggaattctacggcaagttcaaagaaggagttgcatctggcaatctgaacacgatgtttgagtatacctttgattaccagatgaca<br>cctaccaaagtgaaagtccacatgaagaaagcgttatcc |

| | | |
|---|---|---|
| pAG1142 | **MYC**-FRB-$^{FIRE}$mate-IRES-**HA**-mTurquoise2 | atg**gaacaaaagcttatttctgaagaggacttg**gaattcgagatgtggcatgaaggcctggaagaggcatctcgtttgtactttggggaaa<br>ggaacgtgaaaggcatgtttgaggtgctggagcccttgcatgctatgatggaacggggcccccagactctgaaggaaacatcctttaatca<br>ggcctatggtcgagatttaatggaggcccaagagtggtgcaggaagtacatgaaatcagggaatgtcaaggacctcacccaagcctgggac<br>ctctattatcatgtgttccgacgaatctcaaagcaggtctccggaggaggcggcagcggcggaggggatccatggagcatgttgcctttg<br>gcagtgaggacatcgagaacactctggccaatatggacgacgaacaactggataggttggcctttggcgtaattcagctcgatggtgacgg<br>gaatatcctgctgtacaatgctgctgaaggggacatcactggcagagatcccaaacaggtgattgggaagaacttcttcaaggatgttgca<br>cctggaacggatactcccgagttttacggcaaattcaaggaaggcgcagcgtcagggaatctgaacaccatgttcgaatggacgataccga<br>caagcaggggaccaaccaaggtcaaggtgcacttgaagaaagccctttcctaacctcgaggactacaaggacgacgacgacaagcccggga<br>tccgcccctctccctcccccccccctaacgttactggccgaagccgcttggaataaggccggtgtgcgtttgtctatatgttattttccac<br>catattgccgtcttttggcaatgtgagggcccggaaacctggccctgtcttcttgacgagcattcctaggggtctttcccctctcgccaaa<br>ggaatgcaaggtctgttgaatgtcgtgaaggaagcagttcctctggaagcttcttgaagacaaacaacgtctgtagcgaccctttgcaggc<br>agcggaacccccacctggcgacaggtgcctctgcggccaaaagccacgtgtataagatacacctgcaaaggcggcacaaccccagtgcca<br>cgttgtgagttggatagttgtggaaagagtcaaatggctctcctcaagcgtattcaacaaggggctgaaggatgcccagaaggtaccccat<br>tgtatgggatctgatctggggcctcggtgcacatgctttacatgtgtttagtcgaggttaaaaaaacgtctaggccccccgaaccacgggg<br>acgtggttttcctttgaaaaacacgatgataatatggccacaaccatgcgatcg**tacccatacgatgttccagattacgct**gaattcatgg<br>tgagcaagggcgaggagctgttcaccggggtggtgcccatcctggtcgagctggacggcgacgtaaacggccacaagttcagcgtgtccgg<br>cgagggcgagggcgatgccacctacggcaagctgaccctgaagttcatctgcaccaccggcaagctgcccgtgccctggcccaccctcgtg<br>accaccctgtcctggggcgtgcagtgcttcgcccgctaccccgaccacatgaagcagcacgacttcttcaagtccgccatgcccgaaggct<br>acgtccaggagcgcaccatcttcttcaaggacgacggcaactacaagacccgcgccgaggtgaagttcgagggcgacaccctggtgaaccg<br>catcgagctgaagggcatcgacttcaaggaggacggcaacatcctggggcacaagctggagtacaactacttcagcgacaacgtctatatc<br>accgccgacaagcagaagaacggcatcaaggccaacttcaagatccgccacaacatcgaggacggcggcgtgcagctcgccgaccactacc<br>agcagaacacccccatcggcgacggccccgtgctgctgcccgacaaccactacctgagcacccagtccaagctgagcaaagaccccaacga<br>gaagcgcgatcacatggtcctgctggagttcgtgaccgccgccgggatcactctcggcatggacgagctgtacaagtaa |
| pAG1151 | **MYC**-FKBP-$^{FIRE}$tag-IRES-**HA**-IRFP670 | atg**gaacaaaagcttatttctgaagaggacttg**gaattcggagtgcaggtggaaaccatctccccaggagacgggcgcaccttccccaagc<br>gcggccagacctgcgtggtgcactacaccgggatgcttgaagatggaaagaaatttgattcctcccgggacagaaacaagccctttaagtt<br>tatgctaggcaagcaggaggtgatccgaggctgggaagaaggggttgcccagatgagtgtgggtcagagagccaaactgactatatctcca<br>gattatgcctatggtgccactgggcacccaggcatcatcccaccacatgccactctcgtcttcgatgtggagcttctaaaactggaagaat<br>ccggaggaggcggcagcggcggaggggatccggtgacagatattgggtctttgtgaaacgggtgtaactcgaggactacaaggacgacga<br>cgacaagcccgggatccgcccctctccctcccccccccctaacgttactggccgaagccgcttggaataaggccggtgtgcgtttgtctat<br>atgttattttccaccatattgccgtcttttggcaatgtgagggcccggaaacctggccctgtcttcttgacgagcattcctaggggtcttt<br>cccctctcgccaaaggaatgcaaggtctgttgaatgtcgtgaaggaagcagttcctctggaagcttcttgaagacaaacaacgtctgtagc<br>gaccctttgcaggcagcggaacccccacctggcgacaggtgcctctgcggccaaaagccacgtgtataagatacacctgcaaaggcggca<br>caaccccagtgccacgttgtgagttggatagttgtggaaagagtcaaatggctctcctcaagcgtattcaacaaggggctgaaggatgccc<br>agaaggtaccccattgtatgggatctgatctggggcctcggtgcacatgctttacatgtgtttagtcgaggttaaaaaaacgtctaggccc<br>cccgaaccacggggacgtggttttcctttgaaaaacacgatgataatatggccacaaccatgcgatcg**tacccatacgatgttccagatta**<br>**cgct**gaattcatggcgcgtaaggtcgatctcacctcctgcgatcgcgagccgatccacatccccggcagcattcagccgtgcggctgcctg<br>ctagcctgcgacgcgcaggcggtgcggatcacgcgcattacggaaaatgccggcgcgttctttggacgcgaaactccgcgggtcggtgagc<br>tactcgccgattacttcggcgagaccgaagcccatgcgctgcgcaacgcactggcgcagtcttccgatccaaagcgaccggcgctgatctt<br>cggttggcgcgacggcctgaccggccgcaccttcgacatctcactgcatcgccatgacggtacatcgatcatcgagttcgagcctgcggcg<br>gccgaacaggccgacaatccgctgcggctgacgcggcagatcatcgcgcgcaccaaagaactgaagtcgctcgaagagatggccgcacggg<br>tgccgcgctatctgcaggcgatgctcggctatcaccgcgtgatgttgtaccgcttcgcggacgacggctccgggatggtgatcggcgaggc<br>gaagcgcagcgacctcgagagctttctcggtcagcactttccggcgtcgctggtcccgcagcaggcgcggctactgtacttgaagaacgcg<br>atccgcgtggtctcggattcgcgcggcatcagcagccggatcgtgcccgagcacgacgcctccggcgccgcgctcgatctgtcgttcgcgc<br>acctgcgcagcatctcgccctgccatctcgaatttctgcggaacatgggcgtcagcgcctcgatgtcgctgtcgatcatcattgacggcac<br>gctatggggattgatcatctgtcatcattacgagccgcgtgccgtgccgatggcgcagcgcgtcgcggccgaaatgttcgccgacttctta<br>tcgctgcacttcaccgccgcccaccaccaacgctaa |
| pAG1209 | $^{FIRE}$tag-mCherry | atgggtgacagatattgggtctttgtgaaacgggtgagcggcgggggaggctccggagggatggtgagcaagggcgaggaggataacatgg<br>ccatcatcaaggagttcatgcgcttcaaggtgcacatggagggctccgtgaacggccacgagttcgagatcgagggcgagggcgagggccg<br>cccctacgagggcacccagaccgccaagctgaaggtgaccaagggtggccccctgcccttcgcctgggacatcctgtcccctcagttcatg<br>tacggctccaaggcctacgtgaagcaccccgccgacatccccgactacttgaagctgtccttccccgagggcttcaagtgggagcgcgtga<br>tgaacttcgaggacggcggcgtggtgaccgtgacccaggactcctccctgcaggacggcgagttcatctacaaggtgaagctgcgcggcac<br>caacttcccctccgacggccccgtaatgcagaagaagaccatgggctgggaggcctcctccgagcggatgtaccccgaggacggcgccctg<br>aagggcgagatcaagcagaggctgaagctgaaggacggcggccactacgacgctgaggtcaagaccacctacaaggccaagaagcccgtgc<br>agctgcccggcgcctacaacgtcaacatcaagttggacatcacctcccacaacgaggactacaccatcgtggaacagtacgaacgcgccga<br>gggccgccactccaccggcggcatggacgagctgtacaag |
| pAG1241 | mCherry-$^{FIRE}$tag-mCherry | atggtgagcaagggcgaggaggataacatggccatcatcaaggagttcatgcgcttcaaggtgcacatggagggctccgtgaacggccacg<br>agttcgagatcgagggcgagggcgagggccgcccctacgagggcacccagaccgccaagctgaaggtgaccaagggtggccccctgccctt<br>cgcctgggacatcctgtcccctcagttcatgtacggctccaaggcctacgtgaagcaccccgccgacatccccgactacttgaagctgtcc<br>ttccccgagggcttcaagtgggagcgcgtgatgaacttcgaggacggcggcgtggtgaccgtgacccaggactcctccctgcaggacggcg<br>agttcatctacaaggtgaagctgcgcggcaccaacttcccctccgacggccccgtaatgcagaagaagaccatgggctgggaggcctcctc<br>cgagcggatgtaccccgaggacggcgccctgaagggcgagatcaagcagaggctgaagctgaaggacggcggccactacgacgctgaggtc<br>aagaccacctacaaggccaagaagcccgtgcagctgcccggcgcctacaacgtcaacatcaagttggacatcacctcccacaacgaggact<br>acaccatcgtggaacagtacgaacgcgccgagggccgccactccaccggcggcatggacgagctgtacaagtctagaggcggcggctccgg<br>tgacagatattgggtctttgtgaaacgggtgagcggcgggggaggctccggagggatggtgagcaagggcgaggaggataacatggccatc<br>atcaaggagttcatgcgcttcaaggtgcacatggagggctccgtgaacggccacgagttcgagatcgagggcgagggcgagggccgcccct<br>acgagggcacccagaccgccaagctgaaggtgaccaagggtggccccctgcccttcgcctgggacatcctgtcccctcagttcatgtacgg<br>ctccaaggcctacgtgaagcaccccgccgacatccccgactacttgaagctgtccttccccgagggcttcaagtgggagcgcgtgatgaac<br>ttcgaggacggcggcgtggtgaccgtgacccaggactcctccctgcaggacggcgagttcatctacaaggtgaagctgcgcggcaccaact<br>tcccctccgacggccccgtaatgcagaagaagaccatgggctgggaggcctcctccgagcggatgtaccccgaggacggcgccctgaaggg<br>cgagatcaagcagaggctgaagctgaaggacggcggccactacgacgctgaggtcaagaccacctacaaggccaagaagcccgtgcagctg<br>cccggcgcctacaacgtcaacatcaagttggacatcacctcccacaacgaggactacaccatcgtggaacagtacgaacgcgccgagggcc<br>gccactccaccggcggcatggacgagctgtacaag |
| pAG1192 | TOM20(1-34)-ECFP-$^{FIRE}$mate | atggtgggtcggaacagcgccatcgccgcgggcgtgtgcggtgccctcttcatagggtactgcatctactttgaccgcaaaagacgaagtg<br>accccaacttcggatccagtgctggtggtagtgctggtggtatggtgagcaagggcgaggagctgttcaccggggtggtgcccatcctggt<br>cgagctggacggcgacgtaaacggccacaagttcagcgtgtccggcgagggcgagggcgatgccacctacggcaagctgaccctgaagttc<br>atctgcaccaccggcaagctgcccgtgccctggcccaccctcgtgaccaccctgaccgccggcgtgcagtgcttcagccgctaccccgacc<br>acatgaagcagcacgacttcttcaagtccgccatgcccgaaggctacgtccaggagcgcaccatcttcttcaaggacgacggcaactacaa<br>gacccgcgccgaggtgaagttcgagggcgacaccctggtgaaccgcatcgagctgaagggcatcgacttcaaggaggacggcaacatcctg<br>gggcacaagctggagtacaactacatcagccacaacgtctatatcaccgccgacaagcagaagaacggcatcaaggccaacttcaagatcc<br>gccacaacatcgaggacggcagcgtgcagctcgccgaccactaccagcagaacacccccatcggcgacggccccgtgctgctgcccgacaa<br>ccactacctgagcacccagtccgccctgagcaaagaccccaacgagaagcgcgatcacatggtcctgctggagttcgtgaccgccgccggg<br>atcactctcggcatggacgagctgtacaagggagcaagtggaatggagcatgttgcctttggcagtgaggacatcgagaacactctggcca<br>atatggacgacgaacaactggataggttggcctttggcgtaattcagctcgatggtgacgggaatatcctgctgtacaatgctgctgaagg<br>ggacatcactggcagagatcccaaacaggtgattgggaagaacttcttcaaggatgttgcacctggaacggatactcccgagttttacggc<br>aaattcaaggaaggcgcagcgtcagggaatctgaacaccatgttcgaatggacgataccgacaagcaggggaccaaccaaggtcaaggtgc<br>acttgaagaaagccctttcc |

| | | |
|---|---|---|
| pAG1211 | FRB-ECFP-Giantin(3131-3259) | atggcttctagaatcctctggcatgagatgtggcatgaaggcctggaagaggcatctcgtttgtactttggggaaaggaacgtgaaaggca<br>tgtttgaggtgctggagcccttgcatgctatgatggaacggggccccagactctgaaggaaacatcctttaatcaggcctatggtcgaga<br>tttaatggaggcccaagagtggtgcaggaagtacatgaaatcagggaatgtcaaggacctcctccaagcctgggacctctattatcatgtg<br>ttccgacgaatctcaaagactagttatccgtacgacgtaccagactacgcaagtgctggtagtgctggtagtgctggtagtgctggtagtg<br>ctggtagtgctggttcaccggtcgccaccatggtgagcaagggcgaggagctgttcaccggggtggtgcccatcctggtcgagctggacgg<br>cgacgtaaacggccacaagttcagcgtgtccggcgagggcgagggcgatgccacctacggcaagctgaccctgaagttcatctgcaccacc<br>ggcaagctgcccgtgccctggcccaccctcgtgaccaccctgacctggggcgtgcagtgcttcagccgctaccccgaccacatgaagcagc<br>acgacttcttcaagtccgccatgcccgaaggctacgtccaggagcgcaccatcttcttcaaggacgacggcaactacaagacccgcgccga<br>ggtgaagttcgagggcgacaccctggtgaaccgcatcgagctgaagggcatcgacttcaaggaggacggcaacatcctggggcacaagctg<br>gagtacaactacatcagccacaacgtctatatcaccgccgacaagcagaagaacggcatcaaggccaacttcaagatccgccacaacatcg<br>aggacggcagcgtgcagctcgccgaccactaccagcagaacacccccatcggcgacggccccgtgctgctgcccgacaaccactacctgag<br>cacccagtccgccctgagcaaagaccccaacgagaagcgcgatcacatggtcctgctggagttcgtgaccgccgccgggatcactctcggc<br>atggacgagctgtacaagtccggactcagatctcgaggagaaccgcagcaaagcttttctgaagctcagcagcagctatgcaacaccagac<br>aggaagtgaatgaattaaggaagctgctggaagaagaacgagaccaaagagtggctgctgagaatgctctctctgtggccgaggagcagat<br>cagacggttagagcacagtgaatgggactcttcccggactcctatcattggctcctgtggcactcaggagcaggcactgttaatagatctt<br>acaagcaacagttgtcgaaggacccggagtggcgttggatggaagcgagtcctgcgttcactctgtcattcacggacccgagtgccacttc<br>tagcagccatctactttctaatgattcatgtcctgctcattctgtgttttacgggccatcta |
| pAG1212 | mCherry- FKBP-$^{FIRE}$tag | atggtgagcaagggcgaggaggataacatggccatcatcaaggagttcatgcgcttcaaggtgcacatggagggctccgtgaacggccacg<br>agttcgagatcgagggcgagggcgagggccgcccctacgagggcacccagaccgccaagctgaaggtgaccaagggtggccccctgccctt<br>cgcctgggacatcctgtcccctcagttcatgtacggctccaaggcctacgtgaagcaccccgccgacatccccgactacttgaagctgtcc<br>ttccccgagggcttcaagtgggagcgcgtgatgaacttcgaggacggcggcgtggtgaccgtgacccaggactcctccctgcaggacggcg<br>agttcatctacaaggtgaagctgcgcggcaccaacttcccctccgacggccccgtaatgcagaagaagaccatgggctgggaggcctcctc<br>cgagcggatgtaccccgaggacggcgccctgaagggcgagatcaagcagaggctgaagctgaaggacggcggccactacgacgctgaggtc<br>aagaccacctacaaggccaagaagcccgtgcagctgcccggcgcctacaacgtcaacatcaagttggacatcacctcccacaacgaggact<br>acaccatcgtggaacagtacgaacgcgccgagggccgccactccaccggcggcatggacgagctgtacaagtctagaggcggcggctccat<br>ggaacaaaagcttatttctgaagaggacttggaattcggagtgcaggtggaaaccatctccccaggagacgggcgcaccttccccaagcgc<br>ggccagacctgcgtggtgcactacaccgggatgcttgaagatggaaagaaatttgattcctcccgggacagaaacaagccctttaagttta<br>tgctaggcaagcaggaggtgatccgaggctgggaagaaggggttgcccagatgagtgtgggtcagagagccaaactgactatatctccaga<br>ttatgcctatggtgccactgggcacccaggcatcatcccaccacatgccactctcgtcttcgatgtggagcttctaaaactggaagaatct<br>agaggcggcggctccggtgacagatattgggtctttgtgaaacgggtg |
| pAG1141 | **MYC**-FKBP-caged$^{FIRE}$tag-IRES-**HA**-IRFP670 | atg**gaacaaaagcttatttctgaagaggacttg**gaattcggagtgcaggtggaaaccatctccccaggagacgggcgcaccttccccaagc<br>gcggccagacctgcgtggtgcactacaccgggatgcttgaagatggaaagaaatttgattcctcccgggacagaaacaagccctttaagtt<br>tatgctaggcaagcaggaggtgatccgaggctgggaagaaggggttgcccagatgagtgtgggtcagagagccaaactgactatatctcca<br>gattatgcctatggtgccactgggcacccaggcatcatcccaccacatgccactctcgtcttcgatgtggagcttctaaaactggaagaat<br>ccggaggaggcggcagcggcggaggggggatccggtgacagatattgggtctttgtgaaacgggtgggctccggcggcgatgaggtggatgg<br>ctccggcggcgaacatgttgcctttggttcggaagatatcgagaacactctggcgaaaatggacgacggtcaactggatggcttggctttt<br>ggtgccattcaactggatggcgatggcaatatcctccagtataatgcggctgaaggggacattaccggacgtgatccgaaacaggtcattg<br>ggaagaacttcttcaaagacgtagcaccaggtacggatagtccggaattctacggcaagttcaaagaaggagttgcatctggcaatctgaa<br>cacgatgtttgagtatacctttgattaccagatgacacctaccaaagtgaaagtccacatgaagaaagcgttatcctaactcgaggactac<br>aaggacgacgacgacaagcccgggatccgcccctctccctcccccccccctaacgttactggccgaagccgcttggaataaggccggtgtg<br>cgtttgtctatatgttattttccaccatattgccgtcttttggcaatgtgagggcccggaaacctggccctgtcttcttgacgagcattcc<br>taggggtctttcccctctcgccaaaggaatgcaaggtctgttgaatgtcgtgaaggaagcagttcctctggaagcttcttgaagacaaaca<br>acgtctgtagcgaccctttgcaggcagcggaaccccccacctggcgacaggtgcctctgcggccaaaagccacgtgtataagatacacctg<br>caaaggcggcacaaccccagtgccacgttgtgagttggatagttgtggaaagagtcaaatggctctcctcaagcgtattcaacaaggggct<br>gaaggatgcccagaaggtaccccattgtatgggatctgatctggggcctcggtgcacatgctttacatgtgtttagtcgaggttaaaaaaa<br>cgtctaggccccccgaaccacggggacgtggttttcctttgaaaaacacgatgataatatggccacaaccatgcgatcg**tacccatacgat**<br>**gttccagattacgct**gaattcatggcgcgtaaggtcgatctcacctcctgcgatcgcgagccgatccacatccccggcagcattcagccgt<br>gcggctgcctgctagcctgcgacgcgcaggcggtgcggatcacgcgcattacggaaaatgccggcgcgttctttggacgcgaaactccgcg<br>ggtcggtgagctactcgccgattacttcggcgagaccgaagcccatgcgctgcgcaacgcactggcgcagtcttccgatccaaagcgaccg<br>gcgctgatcttcggttggcgcgacggcctgaccggccgcaccttcgacatctcactgcatcgccatgacggtacatcgatcatcgagttcg<br>agcctgcggcggccgaacaggccgacaatccgctgcggctgacgcggcagatcatcgcgcgcaccaaagaactgaagtcgctcgaagagat<br>ggccgcacgggtgccgcgctatctgcaggcgatgctcggctatcaccgcgtgatgttgtaccgcttcgcggacgacggctccgggatggtg<br>atcggcgaggcgaagcgcagcgacctcgagagctttctcggtcagcactttccggcgtcgctggtcccgcagcaggcgcggctactgtact<br>tgaagaacgcgatccgcgtggtctcggattcgcgcggcatcagcagccggatcgtgcccgagcacgacgcctccggcgccgcgctcgatct<br>gtcgttcgcgcacctgcgcagcatctcgccctgccatctcgaatttctgcggaacatgggcgtcagcgcctcgatgtcgctgtcgatcatc<br>attgacggcacgctatggggattgatcatctgtcatcattacgagccgcgtgccgtgccgatggcgcagcgcgtcgcggccgaaatgttcg<br>ccgacttcttatcgctgcacttcaccgccgcccaccaccaacgctaa |
| pFP5290 | ss-$^{FIRE}$mate-**HA**-KDEL-IVS-IRES-ss-**SBP**-mApple-$^{FIRE}$tag-GPI | ATGGATGTATGCGTCCGTCTTGCCCTGTGGCTCCTCTGGGGACTCCTCCTGCACCAGGGCCAGAGCCTCAGCCATAGTCA<br>CAGTGAGAAGGCGACAGGAACCAGCTCGGGCGCCggtttaaacgagcatgttgcctttggcagtgaggacatcgagaaca<br>ctctggccaatatggacgacgaacaactggataggttggcctttggcgtaattcagctcgatggtgacgggaatatcctg<br>ctgtacaatgctgctgaaggggacatcactggcagagatcccaaacaggtgattgggaagaacttcttcaaggatgttgc<br>acctggaacggatactcccgagttttacggcaaattcaaggaaggcgcagcgtcagggaatctgaacaccatgttcgaat<br>ggacgataccgacaagcaggggaccaaccaaggtcaaggtgcacttgaagaaagccctttccggaggtggcggaagt**TAT**<br>**CCtTAtGACGTACCAGACTACGCA**ctcggaccgAAAGATGAACTGTGAgcggccGCATAGATAACTGATCCAGTGTGCTG<br>GAATTAATTCGCTGTCTGCGAGGGCCAGCTGTTGGGGTGAGTACTCCCTCTCAAAAGCGGGCATGACTTCTGCGCTAAGA<br>TTGTCAGTTTCCAAAAACGAGGAGGATTTGATATTCACCTGGCCCGCGGTGATGCCTTTGAGGGTGGCCGCGTCCATCTG<br>GTCAGAAAAGACAATCTTTTTGTTGTCAAGCTTGAGGTGTGGCAGGCTTGAGATCTGGCCATACACTTGAGTGACAATGA<br>CATCCACTTTGCCTTTCTCTCCACAGGTGTCCACTCCCAGGTCCAACTGCAGGTCGAGCATGCATCTAGGGCGGCCAATT<br>CCGCCCCTCTCCCTCCCCCCCCCCTAACGTTACTGGCCGAAGCCGCTTGGAATAAGGCCGGTGTGCGTTTGTCTATATGT<br>TATTTTCCACCATATTGCCGTCTTTTGGCAATGTGAGGGCCCGGAAACCTGGCCCTGTCTTCTTGACGAGCATTCCTAGG<br>GGTCTTTCCCCTCTCGCCAAAGGAATGCAAGGTCTGTTGAATGTCGTGAAGGAAGCAGTTCCTCTGGAAGCTTCTTGAAG<br>ACAAACAACGTCTGTAGCGACCCTTTGCAGGCAGCGGAACCCCCCACCTGGCGACAGGTGCCTCTGCGGCCAAAAGCCAC<br>GTGTATAAGATACACCTGCAAAGGCGGCACAACCCCAGTGCCACGTTGTGAGTTGGATAGTTGTGGAAAGAGTCAAATGG<br>CTCTCCTCAAGCGTATTCAACAAGGGGCTGAAGGATGCCCAGAAGGTACCCCATTGTATGGGATCTGATCTGGGGCCTCG<br>GTGCACATGCTTTACATGTGTTTAGTCGAGGTTAAAAAACGTCTAGGCCCCCCGAACCACGGGGACGTGGTTTTCCTTT<br>GAAAAACACGATGATAAGCTTGCCACAACccgggaGGcgcgccATGTACAGGATGCAACTCCTGTCTTGCATTGCACTAA<br>GTCTTGCACTTGTCACGaattcc**GACGAGAAGACCACTGGTTGGCGAGGTGGACACGTTGTTGAAGGACTGGCTGGGGAA**<br>**CTTGAACAACTTCGTGCACGACTGGAGCATCACCCACAAGGTCAACGTGAACCA**cctgcaggtGTGAGCAAGGGCGAGGA<br>GAATAACATGGCCATCATCAAGGAGTTCATGCGCTTCAAGGTGCACATGGAGGGCTCCGTGAACGGCCACGAGTTCGAGA<br>TCGAGGGCGAGGGCGAGGGCCGCCCCTACGAGGCCTTTCAGACCGCTAAGCTGAAGGTGACCAAGGGTGGCCCCCTGCCC<br>TTCGCCTGGGACATCCTGTCCCCTCAGTTCATGTACGGCTCCAAGGTCTACATTAAGCACCCAGCCGACATCCCCGACTA<br>CTTCAAGCTGTCCTTCCCCGAGGGCTTCAGGTGGGAGCGCGTGATGAACTTCGAGGACGGCGGCATTATTCACGTTAACC<br>AGGACTCCTCCCTGCAGGACGGCGTGTTCATCTACAAGGTGAAGCTGCGCGGCACCAACTTCCCCTCCGACGGCCCCGTA<br>ATGCAGAAGAAGACCATGGGCTGGGAGGCCTCCGAGGAGCGGATGTACCCCGAGGACGGCGCCCTGAAGAGCGAGATCAA<br>GAAGAGGCTGAAGCTGAAGGACGGCGGCCACTACGCCGCCGAGGTCAAGACCACCTACAAGGCCAAGAAGCCCGTGCAGC<br>TGCCCGGCGCCTACATCGTCGACATCAAGTTGGACATCGTGTCCCACAACGAGGACTACACCATCGTGGAACAGTACGAA<br>CGCGCCGAGGGCCGCCACTCCACCGGCGGCATGGACGAGCTGTACAAGggccggCCaggcggcggctccggtgacagata<br>ttgggtctttgtgaaacgggtgggaggtggatcaCGTACGCTGGAAAATGGCGGAACCTCTCTGTCTGAAAAAACTGTGC<br>TGCTGCTGGTGACTCCCTTTCTGGCCGCTGCTTGGTCCCTCCACCCATGA |

| | | |
|---|---|---|
| pFP5299 | ss-$^{FIRE}$mate-**HA**-KDEL-IVS-IRES-ManII-**SBP**-mApple-$^{FIRE}$tag | ATGGATGTATGCGTCCGTCTTGCCCTGTGGCTCCTCTGGGGACTCCTCCTGCACCAGGGCCAGAGCCTCAGCCATAGTCA<br>CAGTGAGAAGGCGACAGGAACCAGCTCGGGCGCCggtttaaacgagcatgttgcctttggcagtgaggacatcgagaaca<br>ctctggccaatatggacgacgaacaactggataggttggcctttggcgtaattcagctcgatggtgacgggaatatcctg<br>ctgtacaatgctgctgaaggggacatcactggcagagatcccaaacaggtgattgggaagaacttcttcaaggatgttgc<br>acctggaacggatactcccgagttttacggcaaattcaaggaaggcgcagcgtcagggaatctgaacaccatgttcgaat<br>ggacgataccgacaagcaggggaccaaccaaggtcaaggtgcacttgaagaaagccctttccggaggtggcggaagt**TAT**<br>**CCtTAtGACGTACCAGACTACGCA**ctcggaccgAAAGATGAACTGTGAgcggccGCATAGATAACTGATCCAGTGTGCTG<br>GAATTAATTCGCTGTCTGCGAGGGCCAGCTGTTGGGGTGAGTACTCCCTCTCAAAAGCGGGCATGACTTCTGCGCTAAGA<br>TTGTCAGTTTCCAAAAACGAGGAGGATTTGATATTCACCTGGCCCGCGGTGATGCCTTTGAGGGTGGCCGCGTCCATCTG<br>GTCAGAAAAGACAATCTTTTTGTTGTCAAGCTTGAGGTGTGGCAGGCTTGAGATCTGGCCATACACTTGAGTGACAATGA<br>CATCCACTTTGCCTTTCTCTCCACAGGTGTCCACTCCCAGGTCCAACTGCAGGTCGAGCATGCATCTAGGGCGGCCAATT<br>CCGCCCCTCTCCCTCCCCCCCCCCTAACGTTACTGGCCGAAGCCGCTTGGAATAAGGCCGGTGTGCGTTTGTCTATATGT<br>TATTTTCCACCATATTGCCGTCTTTTGGCAATGTGAGGGCCCGGAAACCTGGCCCTGTCTTCTTGACGAGCATTCCTAGG<br>GGTCTTTCCCCTCTCGCCAAAGGAATGCAAGGTCTGTTGAATGTCGTGAAGGAAGCAGTTCCTCTGGAAGCTTCTTGAAG<br>ACAAACAACGTCTGTAGCGACCCTTTGCAGGCAGCGGAACCCCCCACCTGGCGACAGGTGCCTCTGCGGCCAAAAGCCAC<br>GTGTATAAGATACACCTGCAAAGGCGGCACAACCCCAGTGCCACGTTGTGAGTTGGATAGTTGTGGAAAGAGTCAAATGG<br>CTCTCCTCAAGCGTATTCAACAAGGGGCTGAAGGATGCCCAGAAGGTACCCCATTGTATGGGATCTGATCTGGGGCCTCG<br>GTGCACATGCTTTACATGTGTTTAGTCGAGGTTAAAAAACGTCTAGGCCCCCCGAACCACGGGGACGTGGTTTTCCTTT<br>GAAAACACGATGATAAGCTTGCCACAACccgggaGGcgcgccATGAAGTTAAGTCGCCAGTTCACCGTGTTTGGCAGCG<br>CGATCTTCTGCGTCGTAATCTTCTCACTCTACCTGATGCTGGACAGGGGTCACTTGGACTACCCTCGGGGCCCGCGCCAG<br>GAGGGCTCCTTTCCGCAGGGCCAGCTTTCAATATTGCAAGAAAAGATTGACCATTTGGAGCGTTTGCTCGCTGAGAACAA<br>CGAGATTATCTCAAATATCAGAGACTCAGTCATCAACCTGAGCGAGTCTGTGGAGGACGGCCCGCGGGGGTCACCAGGCA<br>ACGCCAGCCAAGGCTCCATCCACCTCCACTCGCCACAGTTGGCCCTGCAGGCTGACCCCAGAGACTGTTTGGATCCCACC<br>GGTCGCCACCGGaattcc**GACGAGAAGACCACTGGTTGGCGAGGTGGACACGTTGTTGAAGGACTGGCTGGGGAACTTGA**<br>**ACAACTTCGTGCACGACTGGAGCATCACCCACAAGGTCAACGTGAACCA**cctgcaggtGTGAGCAAGGGCGAGGAGAATA<br>ACATGGCCATCATCAAGGAGTTCATGCGCTTCAAGGTGCACATGGAGGGCTCCGTGAACGGCCACGAGTTCGAGATCGAG<br>GGCGAGGGCGAGGGCCGCCCCTACGAGGCCTTTCAGACCGCTAAGCTGAAGGTGACCAAGGGTGGCCCCCTGCCCTTCGC<br>CTGGGACATCCTGTCCCCTCAGTTCATGTACGGCTCCAAGGTCTACATTAAGCACCCAGCCGACATCCCCGACTACTTCA<br>AGCTGTCCTTCCCCGAGGGCTTCAGGTGGGAGCGCGTGATGAACTTCGAGGACGGCGGCATTATTCACGTTAACCAGGAC<br>TCCTCCCTGCAGGACGGCGTGTTCATCTACAAGGTGAAGCTGCGCGGCACCAACTTCCCCTCCGACGGCCCCGTAATGCA<br>GAAGAAGACCATGGGCTGGGAGGCCTCCGAGGAGCGGATGTACCCCGAGGACGGCGCCCTGAAGAGCGAGATCAAGAAGA<br>GGCTGAAGCTGAAGGACGGCGGCCACTACGCCGCCGAGGTCAAGACCACCTACAAGGCCAAGAAGCCCGTGCAGCTGCCC<br>GGCGCCTACATCGTCGACATCAAGTTGGACATCGTGTCCCACAACGAGGACTACACCATCGTGGAACAGTACGAACGCGC<br>CGAGGGCCGCCACTCCACCGGCGGCATGGACGAGCTGTACAAGggccggCCaggcggcggctccggtgacagatattggg<br>tctttgtgaaacgggtgggcctcgagGGCCaaTTAatTAActctagATAA |
| pFP5370 | ss-$^{FIRE}$mate-**HA**-KDEL-IVS-IRES-TNF-**SBP**-mCherry-$^{FIRE}$tag | ATGGATGTATGCGTCCGTCTTGCCCTGTGGCTCCTCTGGGGACTCCTCCTGCACCAGGGCCAGAGCCTCAGCCATAGTCA<br>CAGTGAGAAGGCGACAGGAACCAGCTCGGGCGCCggtttaaacgagcatgttgcctttggcagtgaggacatcgagaaca<br>ctctggccaatatggacgacgaacaactggataggttggcctttggcgtaattcagctcgatggtgacgggaatatcctg<br>ctgtacaatgctgctgaaggggacatcactggcagagatcccaaacaggtgattgggaagaacttcttcaaggatgttgc<br>acctggaacggatactcccgagttttacggcaaattcaaggaaggcgcagcgtcagggaatctgaacaccatgttcgaat<br>ggacgataccgacaagcaggggaccaaccaaggtcaaggtgcacttgaagaaagccctttccggaggtggcggaagt**TAT**<br>**CCtTAtGACGTACCAGACTACGCA**ctcggaccgAAAGATGAACTGTGAgcggccGCATAGATAACTGATCCAGTGTGCTG<br>GAATTAATTCGCTGTCTGCGAGGGCCAGCTGTTGGGGTGAGTACTCCCTCTCAAAAGCGGGCATGACTTCTGCGCTAAGA<br>TTGTCAGTTTCCAAAAACGAGGAGGATTTGATATTCACCTGGCCCGCGGTGATGCCTTTGAGGGTGGCCGCGTCCATCTG<br>GTCAGAAAAGACAATCTTTTTGTTGTCAAGCTTGAGGTGTGGCAGGCTTGAGATCTGGCCATACACTTGAGTGACAATGA<br>CATCCACTTTGCCTTTCTCTCCACAGGTGTCCACTCCCAGGTCCAACTGCAGGTCGAGCATGCATCTAGGGCGGCCAATT<br>CCGCCCCTCTCCCTCCCCCCCCCCTAACGTTACTGGCCGAAGCCGCTTGGAATAAGGCCGGTGTGCGTTTGTCTATATGT<br>TATTTTCCACCATATTGCCGTCTTTTGGCAATGTGAGGGCCCGGAAACCTGGCCCTGTCTTCTTGACGAGCATTCCTAGG<br>GGTCTTTCCCCTCTCGCCAAAGGAATGCAAGGTCTGTTGAATGTCGTGAAGGAAGCAGTTCCTCTGGAAGCTTCTTGAAG<br>ACAAACAACGTCTGTAGCGACCCTTTGCAGGCAGCGGAACCCCCCACCTGGCGACAGGTGCCTCTGCGGCCAAAAGCCAC<br>GTGTATAAGATACACCTGCAAAGGCGGCACAACCCCAGTGCCACGTTGTGAGTTGGATAGTTGTGGAAAGAGTCAAATGG<br>CTCTCCTCAAGCGTATTCAACAAGGGGCTGAAGGATGCCCAGAAGGTACCCCATTGTATGGGATCTGATCTGGGGCCTCG<br>GTGCACATGCTTTACATGTGTTTAGTCGAGGTTAAAAAACGTCTAGGCCCCCCGAACCACGGGGACGTGGTTTTCCTTT<br>GAAAACACGATGATAAGCTTGCCACAACccgggaGGcgcgccatgtctaccgagtctatgattaggacgtggaactgg<br>ctgaggaggcactgcccaaaaaaccggcggaccacagggctc<br>taggagatgtctgtttctgtctctgttctcttttctcatcgtggctggcgctacaacactcttctgtctgctccatttcg<br>gcgtgattggaccacagcgagaggaatttccccgggatctgtcactcatttcaccactggcacaggctgtccgatcttca<br>tctcggactccatccgacaaacctgtcgcccatgtcgtcgccaacccacaggccgagggccagctccagtggctcaatag<br>gagggcaaacgctctgctcgccaatggcgtggaactccgggataaccagctcgtcgtgcctagtgagggactgtacctca<br>tctactcccaggtgctgtttaagggccagggatgtccttctacacatgtgctgctcacacacacaatttcacggatcgcc<br>gtgtcttaccagactaaagtcaatctgctctctgccatcaaatccccatgtcagcgggaaacacctgagggcgctgaggc<br>taaaccttggtacgaacccatctacctcggaggcgtgttccagctggagaagggcgatagactgagtgctgagatcaatc<br>gacccgactacctcgactttgccgaatctggccaggtctactttggcatcattgctctgggcctcgagggccgagctcat<br>ggcgcacctaggcctttgaattcc**ATGGACGAGAAGACCACTGGTTGGCGAGGTGGACACGTTGTTGAAGGACTGGCTGG**<br>**GGAACTTGAACAACTTCGTGCACGACTGGAGCATCACCCACAAGGTCAACGTGAACCA**<br>cctgcaggtatggtgagcaagggcgaggaggataacatggccatcatcaaggagttcatgcgcttcaaggtgcacatgga<br>gggctccgtgaacggccacgagttcgagatcgagggcgagggcgagggccgcccctacgagggcacccagaccgccaagc<br>tgaaggtgaccaagggtggccccctgcccttcgcctgggacatcctgtcccctcagttcatgtacggctccaaggcctac<br>gtgaagcaccccgccgacatccccgactacttgaagctgtccttccccgagggcttcaagtgggagcgcgtgatgaactt<br>cgaggacggcggcgtggtgaccgtgacccaggactcctccctacaggacggcgagttcatctacaaggtgaagctgcgcg<br>gcaccaacttcccctccgacggccccgtaatgcagaagaagaccatgggctgggaggcctcctccgagcggatgtacccc<br>gaggacggcgccctgaagggcgagatcaagcagaggctgaagctgaaggacggcggccactacgacgctgaggtcaagac<br>cacctacaaggccaagaagcccgtgcagctgcccggcgcctacaacgtcaacatcaagttggacatcacctcccacaacg<br>aggactacaccatcgtggaacagtacgaacgcgccgagggccgccactccaccggcggcatggacgagctgtacaagggc<br>cggccatctagaggcggcggctccggtgacagatattgggtctttgtgaaacgggtgctcgagtga |
| pFP5369 | KIF17MD-flag-$^{FIRE}$mate | atggcctccgaggcggtgaaggttgtcgtgcgctgccgtcccatgaaccagcgggagcgagagctgcgctgccagcccgtggtgactgtg<br>gactgcgcgcgcgcccagtgctgcatccagaacccgggcgccgccgacgagccgcccaagcagttcaccttcgacggcgcctaccacgtg<br>gaccacgtcaccgagcagatctacaacgagatcgcctatccgctggtggagggcgtcactgagggctacaatggcaccatctttgcctac<br>ggccagacaggcagcgggaagtccttcaccatgcagggcctgccggatccgccctcccagagaggcatcatccccagggccttcgagcac<br>gtgttcgagagcgtccagtgtgcagagaacactaagttcctggtccgggcctcctacctggagatctacaatgaagatgtccgggacctc<br>cttggggctgacaccaagcagaagctggagctgaaggagcacccagagaagggcgtgtacgtgaaggggctgtccatgcacacggtgcac<br>agcgtggcccagtgtgagcacatcatggagactggctggaagaaccgttcggtcggctacacgctgatgaacaaggattcctcacgctcg<br>cactccatcttcaccatcagcatcgagatgtctgccgtggatgagcggggcaaggaccacctccgggcgggcaagctgaacctggtggac<br>ctggcgggcagcgagcggcagtccaagaccggggccacgggcgagcggctcaaggaggccaccaagatcaacctgtcgctctcggcactg<br>ggcaatgtcatctcggcgctggtggacgggcgctgtaagcacgtcccctaccgtgactcgaagctgacgcggctgctgcaggactcactg<br>ggcggcaacaccaagacgctcatggtggcctgcctgtcgcctgcggacaacaactacgatgagacactcagcacgctgcgctacgccaac<br>cgggccaagaacatcaggaacaagccgcgcatcaatgaggaccccaaggatgcgctgcttcgcgagtaccaggaggagatcaagaagctc<br>aaggccatcctgacacagcagatgagccccagcagcctgtcagccctgctgtccaggcaggtgcccccagaccctgtgcaggtggaggag<br>aagctgttgccccaacctgtgatccagcatgacgtggaggccgagaagcagctgatccgggaggagtatgaagagcgcctggcccggctg<br>aaagccgactataaggccgagcaggagtctcgggccaggctggaggaagacatcactgccatgcgcaactcatatgacgtcaggctgtcc<br>acgctggaggagaacctgcggaaggagacagaggctgtcctgcaggtgggagtcctctacaaggctgaggtcatgtccagggctgagttt<br>gccagcagcgctgagtacccgcctgcttttcagtatgagacagtggtgaaacccaaggtcttctccacgactgacactctgcccagtgac<br>gatgtctccaagactcaggtttcctccaggtttgcggagctgcccaaggtggaaccctccaaatctgagatttctctgggctccagtgag |

| | | |
|---|---|---|
| | | tcatcctcgctcgaagaaacccttgcggccgctGATTACAAGGATGACGATGACAAGttaattaaagagcatgttgcctttggcagtgag<br>gacatcgagaacactctggccaatatggacgacgaacaactggataggttggcctttggcgtaattcagctcgatggtgacgggaatatc<br>ctgctgtacaatgctgctgaaggggacatcactggcagagatcccaaacaggtgattgggaagaacttcttcaaggatgttgcacctgga<br>acggatactcccgagttttacggcaaattcaaggaaggcgcagcgtcagggaatctgaacaccatgttcgaatggacgataccgacaagc<br>aggggaccaaccaaggtcaaggtgcacttgaagaaagccctttcctga |
| FP5380 | LAMP1-SBP-mCherry-FIREtag | ATGGCGGCCCCCGGCAGCGCCCGGCGACCCCTGCTGCTGCTACTGCTGTTGCTGCTGCTCGGCCTCATGCATTGTGCGTCAGCAGCAATG<br>TTTATGGTGAAAAATGGCAACGGGACCGCGTGCATAATGGCCAACTTCTCTGCTGCCTTCTCAGTGAACTACGACACCAAGAGTGGCCCT<br>AAGAACATGACCTTTGACCTGCCATCAGATGCCACAGTGGTGCTCAACCGCAGCTCCTGTGGAAAAGAGAACACTTCTGACCCCAGTCTC<br>GTGATTGCTTTTGGAAGAGGACATACACTCACTCTCAATTTCACGAGAAATGCAACACGTTACAGCGTCCAGCTCATGAGTTTTGTTTAT<br>AACTTGTCAGACACACACCTTTTCCCCAATGCGAGCTCCAAAGAAATCAAGACTGTGGAATCTATAACTGACATCAGGGCAGATATAGAT<br>AAAAAATACAGATGTGTTAGTGGCACCCAGGTCCACATGAACAACGTGACCGTAACGCTCCATGATGCCACCATCCAGGCGTACCTTTCC<br>AACAGCAGCTTCAGCAGGGGAGAGACACGCTGTGAACAAGACAGGCCTTCCCCAACCACAGCGCCCCCTGCGCCACCCAGCCCCTCGCCC<br>TCACCCGTGCCCAAGAGCCCCTCTGTGGACAAGTACAACGTGAGCGGCACCAACGGGACCTGCCTGCTGGCCAGCATGGGGCTGCAGCTG<br>AACCTCACCTATGAGAGGAAGGACAACACGACGGTGACAAGGCTTCTCAACATCAACCCCAACAAGACCTCGGCCAGCGGGAGCTGCGGC<br>GCCCACCTGGTGACTCTGGAGCTGCACAGCGAGGGCACCACCGTCCTGCTCTTCCAGTTCGGGATGAATGCAAGTTCTAGCCGGTTTTTC<br>CTACAAGGAATCCAGTTGAATACAATTCTTCCTGACGCCAGAGACCCTGCCTTTAAAGCTGCCAACGGCTCCCTGCGAGCGCTGCAGGCC<br>ACAGTCGGCAATTCCTACAAGTGCAACGCGGAGGAGCACGTCCGTGTCACGAAGGCGTTTTCAGTCAATATATTCAAAGTGTGGGTCCAG<br>GCTTTCAAGGTGGAAGGTGGCCAGTTTGGCTCTGTGGAGGAGTGTCTGCTGGACGAGAACAGCATGCTGATCCCCATCGCTGTGGGTGGT<br>GCCCTGGCGGGGCTGGTCCTCATCGTCCTCATCGCCTACCTCGTCGGCAGGAAGAGGAGTCACGCAGGCTACCAGACTATCttgaattcc<br>ATGGACGAGAAGACCACTGGTTGGCGAGGTGGACACGTTGTTGAAGGACTGGCTGGGGAACTTGAACAACTTCGTGCACGACTGGAGCAT<br>CACCCACAAGGTCAACGTGAACCAcctgcaggtATGGTGAGCAAGGGCGAGGAGGATAACATGGCCATCATCAAGGAGTTCATGCGCTTC<br>AAGGTGCACATGGAGGGCTCCGTGAACGGCCACGAGTTCGAGATCGAGGGCGAGGGCGAGGGCCGCCCCTACGAGGGCACCCAGACCGCC<br>AAGCTGAAGGTGACCAAGGGTGGCCCCCTGCCCTTCGCCTGGGACATCCTGTCCCCTCAGTTCATGTACGGCTCCAAGGCCTACGTGAAG<br>CACCCCGCCGACATCCCCGACTACTTGAAGCTGTCCTTCCCCGAGGGCTTCAAGTGGGAGCGCGTGATGAACTTCGAGGACGGCGGCGTG<br>GTGACCGTGACCCAGGACTCCTCCCTaCAGGACGGCGAGTTCATCTACAAGGTGAAGCTGCGCGGCACCAACTTCCCCTCCGACGGCCCC<br>GTAATGCAGAAGAAGACCATGGGCTGGGAGGCCTCCTCCGAGCGGATGTACCCCGAGGACGGCGCCCTGAAGGGCGAGATCAAGCAGAGG<br>CTGAAGCTGAAGGACGGCGGCCACTACGACGCTGAGGTCAAGACCACCTACAAGGCCAAGAAGCCCGTGCAGCTGCCCGGCGCCTACAAC<br>GTCAACATCAAGTTGGACATCACCTCCCACAACGAGGACTACACCATCGTGGAACAGTACGAACGCGCCGAGGGCCGCCACTCCACCGGC<br>GGCATGGACGAGCTGTACAAGggccggCCatctagaggcggcggctccGGTGACAGATATTGGGTCTTTGTGAAACGGGTGCTCGAGTGA |

**Table S2. Experimental conditions**

| Figure | plasmids | Ratio |
|---|---|---|
| Fig. 1c,e,i | pAG1159 + pAG1160 | 5:1 |
| Fig. 1g | pAG1159 + pAG1344 | 5:1 |
| Fig. 1k | pAG1171 + pAG1160 | 5:1 |
| Fig. 1m | pAG1169 + pAG1160 | 5:1 |
| Fig. 1o | pAG1186 + pAG1160 | 5:1 |
| Fig. 1q | pAG1171 + pAG1160 | 5:1 |
| Fig. 2b | pAG1243 + pAG1242 | 5:1 |
| Fig. 2f | pAG1247 + pAG1280 | 5:1 |
| Fig. 3b | pFP5370 | |
| Fig. 3c | pFP5290 | |
| Fig. 3d | pFP5299 | |
| Fig. 4b | pFP5380 + pFP5369 | 5:1 |
| Fig. 5a | pAG1237 + pAG1160 | 5:1 |
| Fig. 5c | pAG1237 + pAG1238 | 5:1 |
| Fig. 6a,d | pAG1247 + pAG1244 | 5:1 |
| Fig. 6h | pAG1247 + pAG1345 | 5:1 |
| Fig. S1 | pAG1142 + pAG1151 | 1:1 |
| Fig. S3a,c | pAG1159 + pAG1160 | 5:1 |
| Fig. S4a | pAG1159 + pAG1209 | 5:1 |
| Fig. S4c | pAG1159 + pAG1241 | 5:1 |
| Fig. S5a | pAG1171 + pAG1160 | 5:1 |
| Fig. S5c | pAG1169 + pAG1160 | 5:1 |
| Fig. S5e | pAG1186 + pAG1160 | 5:1 |
| Fig. S6 | pAG1192 + pAG1211 + pAG1212 | 5:5:1 |
| Fig. S8b | pAG1141 + pAG1142 | 1:1 |
| Fig. S8c | pAG1171 + pAG1188 | 5:1 |
| Fig. S9a | pAG1237 + pAG1160 | 5:1 |
| Fig. S9c | pAG1237 + pAG1238 | 5:1 |
